# Dynamic Modeling, Parameter Estimation and Uncertainty Analysis in 𝗥

**DOI:** 10.1101/085001

**Authors:** Daniel Kaschek, Wolfgang Mader, Mirjam Fehling-Kaschek, Marcus Rosenblatt, Jens Timmer

**Affiliations:** University of Freiburg

**Keywords:** Dynamic models, parameter estimation, code generation, maximum likelihood, uncertainty analysis

## Abstract

In a wide variety of research fields, dynamic modeling is employed as an instrument to learn and understand complex systems. The differential equations involved in this process are usually non-linear and depend on many parameters whose values decide upon the characteristics of the emergent system. The inverse problem, i.e. the inference or estimation of parameter values from observed data, is of interest from two points of view. First, the existence point of view, dealing with the question whether the system is able to reproduce the observed dynamics for any parameter values. Second, the identi ability point of view, investigating invariance of the prediction under change of parameter values, as well as the quanti cation of parameter uncertainty.

In this paper, we present the R package **dMod** providing a framework for dealing with the inverse problem in dynamic systems. The particularity of the approach taken by **dMod** is to provide and propagate accurate derivatives computed from symbolic expressions wherever possible. This derivative information highly supports the convergence of optimization routines and enhances their numerical stability, a requirement for the appli-cability of so sticated uncertainty analysis methods. Computational efficiency is achieved by automatic generation and execution of C code. The framework is object oriented (S3) and provides a variety of functions to set up dynamic models, observation functions and parameter transformations for multi-conditional parameter estimation.

The key elements of the framework and the methodology implemented in **dMod** are highlighted by an application on a three-compartment transporter model.

## 1. Introduction

Dynamic models are found in several research fields, such as physics, biology or finance. In all these fields, models link theoretical concepts and empirical evidence. Single mechanisms or single processes of a complex system are represented by respective terms in the equations of a dynamic model. Parameter estimation can then identify those processes which are crucial to explain the observation. In that sense, parameter estimation can be employed as an instrument to *understand* complex systems. Once, the link between observation and model is established by the estimated parameters, questions about their identi ability arise. The parameter space needs to be explored to analyze whether the estimate is unique and to determine con dence bounds.

Although the problem of parameter estimation in non-linear dynamic systems is highly relevant and at the heart of statistical computing, to this day there are not more than four R packages published on the topic on the Comprehensive R Archive Network (CRAN), namely **FME** (Soetaert and Petzoldt 2010), **nlmeODE** (Tornoe 2012), **mkin** (Ranke, Lindenberger, and Lehmann 2016) and **scaRabee** (Bihorel 2014). All packages have in common that they are built upon the **deSolve** package (Soetaert, Petzoldt, and Setzer 2010). The packages mkin and **FME** support ODEs defined by compiled code where **mkin** also provides tools to autogenerate the C code and compile it using the inline package. For model fitting and uncertainty analysis, **mkin** fully resorts to the functionality of **FME**. Concerning model fitting, **FME** and **nlmeODE** (with **nlme** in the background) use deterministic derivative-based optimizers by default, i.e. either Levenberg-Marquardt or Newton methods provided by `nls.lm(), optim() or nlmimb()`. Although all these optimizers support gradient or Hessian information as input, only the **nlmeODE** package provides an option to augment the ODE by its sensitivity equations to generate derivates for the residuals. By default, sensitivities are computed by finite differences. Last but not least, the **scaRabee** package uses the Nelder-Mead optimization algorithm which is derivative-free but needs in general more iterations until convergence compared to derivative-based methods. In all packages, uncertainty analysis is by default based on the variance-covariance matrix, i.e. on the inverse Hessian matrix of the least squares function. For non-linear models, this method provides a good approximation only if parameters are identi able and the data is highly informative. If these conditions are not met, more sophisticated methods like e.g. Markov chain Monte Carlo (MCMC) sampling, implemented in **FME**, are required. The strength of **scaRabee** and especially **nlmeODE** is multi-conditional fitting. This means that the same model with the same parameters but different forcings to re ect experimental conditions is tted simultaneously to the condition-specific data sets. In the context of mixed-e ects modeling as provided by **nlmeODE**, parameters can be grouped in fixed effects (same parameters over all conditions) and random effects (parameters are different between conditions).

In this paper we present **dMod**, an R package on dynamic modeling and parameter estimation. The aim and core functionality of **dMod** is

i. facilitated set-up of dynamic models with automated C code generation for fast simulation of model predictions and model sensitivities,
ii. exible set-up of general parameter transformations (explicit or implicit) and observation functions, allowing for the implementation of multiple experimental conditions similar to mixed-effects modeling,
iii. parameter estimation based on trust-region optimization of the negative log-likelihood, making use of the sensitivity equations of the dynamic system and of symbolic derivatives of the observation- and parameter transformation functions,
iv. identi ability and uncertainty analysis based on the pro le-likelihood method to deter-mine con dence intervals for parameters and predictions.

The core functionality is extended by two symbolic methods implemented in Python and interfaced via the **rPython** package: identi ability and observability analysis based on Lie-group symmetries (Merkt, Timmer, and Kaschek 2015) and steady-state constraints for parameter estimation (Rosenblatt, Timmer, and Kaschek 2016).

The consequent implementation of capabilities to generate and propagate derivatives on a compiled-code level distinguishes **dMod** from other modeling frameworks as discussed above. Most of the standard optimization routines implemented in R need derivative information. However, the computation of derivatives in the context of ODE models holds some pitfalls. The accuracy of sensitivities obtained from nite differences can be insu cient because the step control of the integrator presents an additional source of numeric inaccuracy. Even for numeric methods circumventing this problem, e.g. complex-step derivatives (Squire and Trapp 1998), the use of sensitivity equations is still bene cial judging from the accuracy vs. computational cost ratio. See (Raue, Schilling, Bachmann, Matteson, Schelker, Kaschek, Hug, Kreutz, Harms, Theis, Klingmüller, and Timmer 2013b) for a comparison of methods. Another distinguishing feature of **dMod** is the handling of non-identi ability of parameters, a phenomenon occurring frequently in the context of parameter estimation in dynamic systems. In some cases, non-identi ability has structural reasons. The differential equations bear certain symmetries and these can or cannot be broken, depending on the structure of the observation. A functional relationship between parameters that leaves the observation invariant is the consequence of the latter. In other cases, the data allows to determine a unique optimum but other solutions, although worse, cannot be statistically rejected. The **dMod** package deals with parameter identi ability and parameter uncertainty by the pro le-likelihood method (Murphy and Van der Vaart 2000; Raue, Kreutz, Maiwald, Bachmann, Schilling, Klingmüller, and Timmer 2009; Kreutz, Raue, Kaschek, and Timmer 2013). The method has proven especially useful in the case of non-identi able parameters where re-sults obtained from both, the quadratic approximation by the variance-covariance matrix and MCMC sampling can be misleading (Raue, Kreutz, Theis, and Timmer 2013a). Besides pa-rameter uncertainties, the pro le likelihood method allows to estimate prediction uncertainty (Kreutz, Raue, and Timmer 2012). It therefore supports the planning of new informative experiments (Raue, Kreutz, Maiwald, Klingmüller, and Timmer 2011) or suggests possible model reductions (Maiwald, Hass, Steiert, Vanlier, Engesser, Raue, Kipkeew, Bock, Kaschek, Kreutz, and Timmer 2016).

The key methods implemented in **dMod** are illustrated in great detail on a dynamic model of bile acid flow. The example is a show-case of how modeling is a dynamic process, using the analysis tools implemented in **dMod** to predict, plan new experiments, combine data from different experiments and include non-linear parameter constraints to improve parameter identi ability. Thereby, we show that **dMod** is a full-grown, flexible modeling environment, fast, thanks to compiled code, reliable, thanks to symbolic derivatives, and accurate, thanks to advanced statistical methods.

The paper is organized as follows: Section 2 introduces the mathematical set-up of dynamic modeling, symmetries in dynamic systems, parameter estimation by the maximum-likelihood method and the pro le likelihood. Section 3 discusses the implementation and design principles behind the **dMod** software. The functionality of **dMod** is presented in Section 4 on the example of bile acid flow in a three-compartment model and finally, Section 5 discusses the two Python extensions shipped with **dMod**.

The **dMod** package is available on the Comprehensive R Archive Network at http://CRAN.R-project.org/package=dMod. The project is hosted on GitHub, https://github.com/dkaschek/dMod, where the most recent development version is available. **dMod** is licensed under the GPL-3 license.

## 2. Theoretical background

### 2.1. Dynamic models and model sensitivities

Dynamic models describe systems with states *x*, usually quantifying the involved species, their interaction and evolution over time. The time evolution of the states is expressed via a set of ODEs, 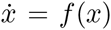. Although constituting quite a special class of dynamic systems, chemical reaction networks formulated by the law of mass action as considered here allow for surprisingly general applications. Typical ODE examples derived from the law of mass action are:

- 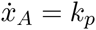, constant production of 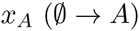
- 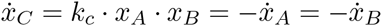, complex formation (A + B → C)
- 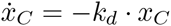, proportional degradation of 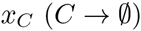

Each generally written as

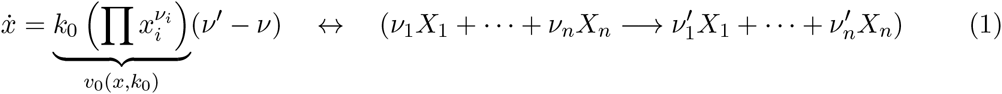

where *X*_1_; …, *X*_*n*_ are the *n* species involved, *ν* and *ν’* encode the stoichiometry of the reaction and *k*_0_ is the reaction rate. When several reactions are involved, eq. (1) generalizes to

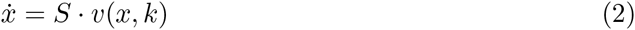

with the stoichiometry matrix *S* obtained from the collection of vectors *v’*–*ν* for reactions 1; …; *r*, and the ux vector 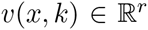. Eq. (2) also holds for general ODEs that do not follow from mass action kinetics. Moreover the system may be explicitly time-dependent and may contain forcings *u(t)* which e.g. describe the external stimulation of the system. A general form of the dynamic model is therefore given by

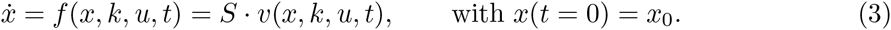

The solution *x*(*t; θ*) for a given parameter vector *θ* = (*k; x*_0_) is called model prediction. It is generally assumed that the stoichiometry matrix *S* of the system is known. Besides the model predition itself, also the sensitivity of the prediction to changes in the parameter values is of interest. The sensitivities 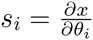 satisfy the *sensitivity equations*

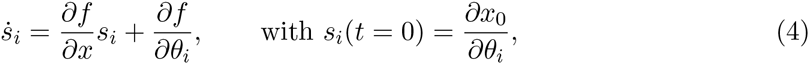

a system of ordinary differential equations that in general depend on the states *x* and, therefore, need to be solved jointly with the original ODE.

Often, the experimentally observed quantities *y* do not directly correspond to the species *x* described by the ODE, but are obtained via an observation function 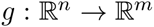,

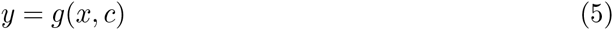

with the observation parameters *c*. Examples for observation functions are scaling and offset transformations, *y* = *c_s_ · x*+*c_o_*, or the measurement of a superposition of species, 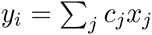. The model prediction for the observed states is obtained by evaluating the observation function on the solution of the ODE. Following the chain rule of differentation also the sensitivities of the observed states 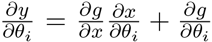 obtained. Here, the parameter vector *θ* has been augmented by the observation parameters, *θ* = (*k; x*_0_; *c*).

The estimation of the parameters given the observation *y*(*t*) is addressed in the next section.

### 2.2. Maximum-likelihood method

Parameter estimation is a common task in statistics. It describes the process of inferring parameter values or parameter ranges of a statistical model based on observed data. Over decades, appropriate estimators have been developed for different problem classes. The principle of maximum likelihood allows to derive an estimator which is especially suited for applications where the distribution of the measurement noise is known. This knowledge about the structure of the noise constitutes additional information that makes the maximum-likelihood estimator (MLE) *efficient*, i.e. from all unbiased estimators the MLE has the lowest variance. Other properties of maximum-likelihood estimation are *consistency*, i.e. the estimated parameter value approaches the true parameter value in the limit of in nite data-sample size, and *asymptotic normality*, i.e. in the limit of in nite data the MLE follows a multivariate normal distribution. See (Azzalini 1996) for an introduction to likelihood theory.

Maximum-likelihood estimation is based on the maximization of the likelihood function *L*(*θ*) = *ϕ*(*y^D^*|*θ*). Here, is the joint probability density for a vector of observations y, c.f. eq. (5), evaluated at the point *y* = *y^D^*, the vector of data points. The distribution *ϕ* depends parametrically on model parameters *θ*. The maximum-likelihood estimator 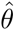 is defined as

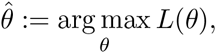

meaning 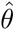 that is an extremum estimator. Depending on the model class and the probability distribution ϕ, maximization of the likelihood can be a challenge beyond the scope of analytical methods. Numerical optimization methods help to solve the maximization problem in practical applications.

The easiest and one of the most frequent situations is when *ϕ* follows a normal distribution, 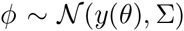. When measurements are statistically independent, the variance-covariance matrix 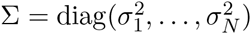 is diagonal and *ϕ* factorizes. The likelihood function reads

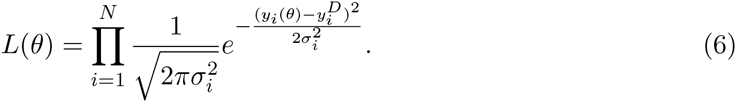

Taking twice the negative logarithm, eq. (6) turns into

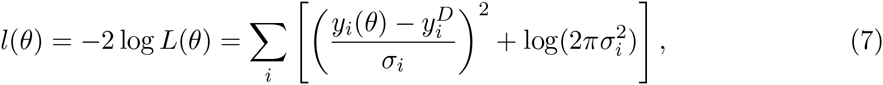

thus converting the maximization of *L*(*θ*) to a minimization of *l*(*θ*). Assuming that the data uncertainties *σ_i_* are known, eq. (7) is the weighted least-squares function shifted by a constant. For unknown *σ_i_*, eq. (7) can be optimized with respect to *θ* and *σ* jointly, yielding the MLE 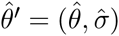

### 2.3. Non-linear optimization

Numerical optimization is a diverse field with as many algorithms as there are optimization problems around. For our application, derivative-based methods, and in particular the Newton method, play a major role.

Optimization by the Newton method attempts to iteratively find the root of the gradient 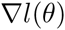 by the recursion

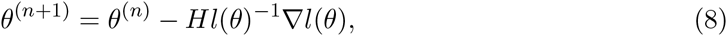

where *Hl*(*θ*) denotes the Hessian, i.e. the matrix of second derivatives of *l*(*θ*). The fact that for normally distributed noise the log-likelihood is a least squares function, has a big advantage: gradient and (approximate) Hessian can be computed from first-order derivatives (Press, Teukolsky, Vetterling, and Flannery 1996). Since the second-order contributions to the Hessian can be neglected, gradient and Hessian read

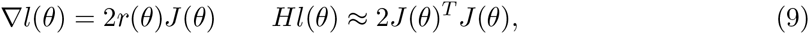

with the weighted residual vector 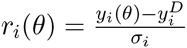 and its first derivative, the Jacobian matrix

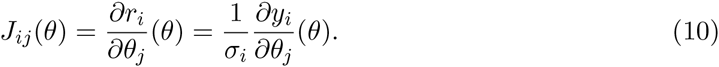

As indicated by eqs. (9)-(10), first order model sensitivities 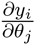 are sufficient to compute gradient and Hessian in good approximation.

The Newton recursion, eq. (8), converges in one iteration if *l*(*θ*) is a quadratic form. However, *l*(*θ*) is only quadratic if the model *y*(*θ*) is linear in which is usually not the case for ODE models. On the other hand, each smooth function, no matter if quadratic or not, can be approximated by a quadratic function based on its Taylor series. Thus, *l*(*θ*) can be approximated by a quadratic function in at least a small region around *θ*. The idea of trust-region optimization is to con ne a Newton step to a region, the trust region, where the quadratic approximation holds (Wright and Nocedal 1999). In each iteration, the trust-region radius is adjusted and the optimization problem restricted to the trust region is solved. For parameter estimation in ODE models this has the additional advantage that parameter changes are constricted and optimization steps into parameter regions where the ODE cannot be numerically solved any more become less frequent. This make trust-region optimization the method of choice for ODEs.

### 2.4. Uncertainty analysis

Parameter estimates 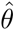 are obtained by non-linear optimization of the log-likelihood function *l*(*θ*). Although being a function of the parameters, the log-likelihood depends on the data, too. Consequently, ‘equivalent’ random realizations of the data would lead to different parameter estimates 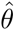. Given 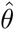 for one random realization of the data, the question is, at which signi cance level we can reject other parameters *θ*. This leads to the related question of parameter convidence intervals.

A useful tool to derive con dence intervals beyond the scope of Fisher Information Matrix is the pro le likelihood. Consider a parameter of interest, *θ_i_*, being one of the parameters in the vector *θ*. Furthermore, let 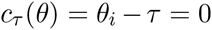 be a parameter constraint. Then, extending the likelihood with the constraint via a Lagrange multiplier λ yields

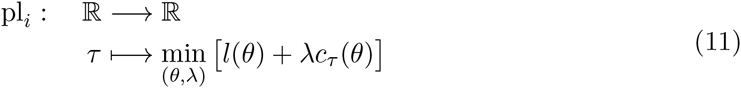

which is called the pro le likelihood of the *i*^th^ parameter. The path 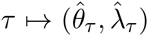 along the minimizing parameters is called the pro le likelihood path. Hence, the pro le likelihood pl_i_ returns the minimal log-likelihood under the constraint 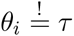. By construction, the inequality 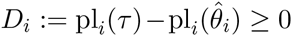 holds for all *τ*, assuming equality at least for 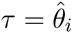. Interpreting the unconstrained model as the null model, which we assume to be true, and that with *θ_i_* fixed to *τ* as the alternative model, the value *D* is twice the log of the likelihood-ratio between those. Hence, a likelihood-ratio test can be performed to accept or reject the alternative model at a given confidence level. To compute the corresponding threshold, we assume Wilks’ theorem according to which the thresholds are the quantiles of the *χ*^2^ distribution with one degree of freedom. Consequently, the 68%/90%/95%-con dence intervals of *θ_i_* are those values of *τ* for which *D_i_*(*τ*) ≤ 1, 2.71 and 3.84, respectively.

The reason to formulate the pro le likelihood by Lagrangian multipliers is that we can directly use eq. (11) to derive the pro le likelihood path 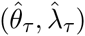 with respect to τ:

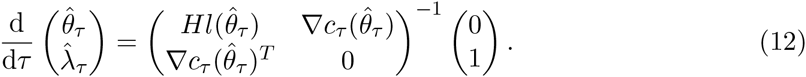

Eq. (12) forms the basis of what is implemented in **dMod** for computing the pro le likelihood. The pro le likelihood approach can be applied to compute con dence intervals for model predictions, too, see (Kreutz *et al*. 2012; Hass, Kreutz, Timmer, and Kaschek 2016).

## 3. Implementation and design principles

The guiding idea behind the implementation of **dMod** is to provide a class structure for models, predictions, observations and parameter transformations that allows a flexible combination of many experimental conditions in one objective function to fit models to data. This flexibility is achieved by two concepts: (1) concatenation of functions by the “*” operator and (2) stacking of functions representing different conditions by the “+” operator. The handling and propagation of derivatives is part of the classes and happens in the background. Methods for the generic `print(), plot() and summary()` functions are implemented for most outputs.

### 3.1. Model formulation

A system of ODEs 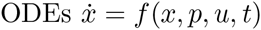 is represented by a named `character` vector of symbolic expressions, the right-hand side of the ODE, involving symbols for the states *x*, the parameters p, the forcings u and the keyword `time` for t. The names of the vector are the state names. The class provided by **dMod** for such objects is the `eqnvec` class which checks whether the character can be parsed and thus, can be interpreted as an equation.

**dMod** provides several ways to define the differential equations. An eqnvec of equations can be explicitly formulated, analogously to a c() command. Especially for chemical reactions it can be tedious to keep track of all gain and loss terms. Therefore, **dMod** provides also the `eqnlist` class which encodes the ODE by a list of (1) state names, (2) rate expressions, (3) compartment volumes and (4) the stoichiometric matrix. Each reaction flux, c.f. eq. (2), occurs only once and the gain and loss is represented by the coe cients in the stoichiometric matrix. **dMod** supports the read-in of a csv file with the stoichiometric matrix and rate expressions and provides a function `addReaction()` to construct an `eqnlist` object step-by-step or add further reactions to an already existing `eqnlist`. An `eqnlist` is converted to (differential) equations, i.e. an eqnvec, by as.eqnvec(). The conversion does all the bookkeeping of gain and loss terms and volume ratios due to compartment transitions.

**dMod** makes use of derivatives wherever possible. It is one of the core functionalities of the **cOde** package upon which **dMod** is based to augment a system of ODEs by its sensitivity equations. If *r* parameters are involved and the system consists of *n* states, the number of equations grows as quick as *n*^2^ + *nr*. Solving the equations can be considerably accelerated by utilizing compiled code. **dMod** provides the `odemodel` class. An object of class `odemodel` is generated from an `eqnlist` or `eqnvec`. In the background, C code for the ODE and the combined system of ODE and sensitivity equations is generated, written to the working directory and compiled. `The odemodel` object keeps track of the shared objects. It is the basis of a prediction function *x*(*t*, *p*) generated by the `Xs()` command.

### 3.2. Prediction functions

**dMod** seeks to stay close to the mathematical formulation, i.e. `x <- Xs(myodemodel)` will indeed return an R function, an object of class `prdfn`, which expects arguments `times` and `pars` and turns them into a model prediction. The letter `“s”` in `“Xs”` refers to sensitivities, i.e. the solution *x*(*t*) is returned, the sensitivities 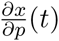 (t) are returned, too.

Besides parameters, the prediction might also depend on forcings *u*(*t*) and sudden events, e.g. setting states to 0 at a predefined point in time. Since forcings and events are by default fixed, they are defined together with the prediction function, `x <- Xs(odemodel, forcings = myforcings, events = myevents)`. The definitions of both, forcings and events, are compatible with the way they are defined in **deSolve**.

### 3.3 Observation functions

An observation function is a function *g*(*x*, *p*_obs_, *t*) that evaluates the solution *x*(*t*) together with additional (observation) parameters *p*_obs_. The function can explicitly depend on time t. Similar to the ODE case, an observation function can be expressed as a character vector with names corresponding to the names of the observables and equations involving symbols for the states, parameters and the keyword time. An observation function is defined as an eqnvec and turned into an `R` function via `Y()`.

It is one of the fundamental concepts of **dMod** to allow concatenation of functions via the “*” operator. The mathematical formulation 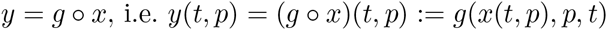 becomes y = g*x in `R`. To obtain derivatives 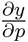, the chain rule is applied: 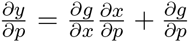 This means that `Y()` needs to be informed which symbols are states and parameters to gen-erate the corresponding expressions 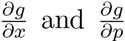. The observation function g takes care of computing theses derivatives from the symbolic expressions and doing the matrix multiplica-tion with the sensitivities 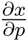 from the prediction function. Evaluation of symbolic expressions can become ine cient in `R`. Therefore, the observation function is usually translated into a `C` code and compiled.

When observation- and prediction functions are concatenated, the result is a prediction function, e.g. `y = g*x` is the `R` function computing values *y*(*t*) from the arguments times and pars via evaluation of the ODE and subsequent evaluation of the observation function. Two observation functions can be concatenated, too, again yielding an observation function.

### 3.4 Parameter transformations

Parameter transformations are the key element of **dMod** to formulate different kinds of constraints and allow the combination of several experimental/modeling conditions in one parameter vector.

In principle, a parameter transformation *p* = Φ(*θ*) is a (differentiable) function connecting inner parameters *p* with **outer** parameters *θ*. The rationale behind the distinction of inner and outer parameters is that the vector *p* usually desribes those parameters defined in the model equations. The outer parameters *θ* refer to a convenient parameterization by which the model parameters are computed. Examples are a log-transform of the inner parameters, 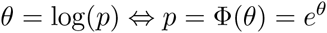 or parameter constraints like 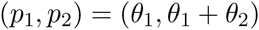.

An `R` function of class parfn is produced by the P() command. Transformations can either be formulated explicitly or implicitly. In the explicit case, the function *p* = Ф(*θ*) corresponds to an eqnvec whose names are the names of the inner parameters and entries are equations with symbols for the outer parameters. An implicit transformation has the form *f*(*p* = Φ(*θ*),*θ*)= 0. In this case, *f* is expressed by an `eqnvec` with equations containing symbols for *p* and *θ* and the names of the eqnvec are the symbols for *p*.

Similar to prediction- and observation functions, parameter functions not only return parameter values but the Jacobian of the transformation, too. Exploiting the chain rule, the derivatives are propagated, allowing to define y = g*x*p which is a function returning *y*(*t;θ*) and 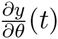, i.e. `g*x*p` is a prediction function.

Let `g, x, p1` and `p2` be an observation-, a prediction- and two parameter transformation functions. Then `p1*p2` is a parameter transformation function, `x*p1` is a prediction function and `g*p1` is an observation function.

### 3.5. Multi-conditional prediction

A set of parameter values, forcings and events capture a certain condition in which we find the modeled system. Manipulating the system, single model parameters, forcings or events need to be changed to account for the manipulation. It is a typical approach in systems inference to systematically perturb small parts of a modeled system to reveal information about the processes.

The aim of **dMod** is to allow for “simultaneous” predictions under several conditions and compare these predictions to the corresponding experimental data sets to estimate model parameters. Different experimental conditions are typically expressed by that fact that some parameters are different between conditions whereas others are common to all conditions. This situation occurs e.g. if perturbation experiments are performed, affecting only few parts of a system. Mathematically speaking, we want to construct a parameter transformation

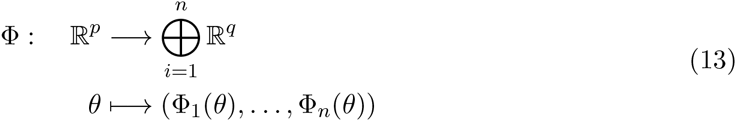

where *i* = 1*, …, n* corresponds to the different conditions. If all parameters are shared throughout all conditions, then *p* = *q*. If, however, all parameters are distinct, then 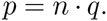 Perturbation experiments correspond to a situation where 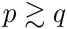.

The **dMod** package allows to define transformations Φ, see eq. (13), by the “+” operator:

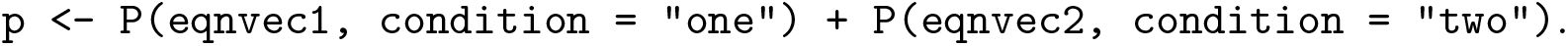

All symbols from `eqnvec1` and `eqnvec2` are collected and their union constitutes the symbols of *θ*. The evaluation `p(theta)` returns a list of length *n* (in the example *n* = 2) with inner parameters. The “+” operator can be applied consecutively to add conditions.

Prediction- and observation functions are generalized to multiple conditions by the “+” operator, too. Mathematically speaking, they become

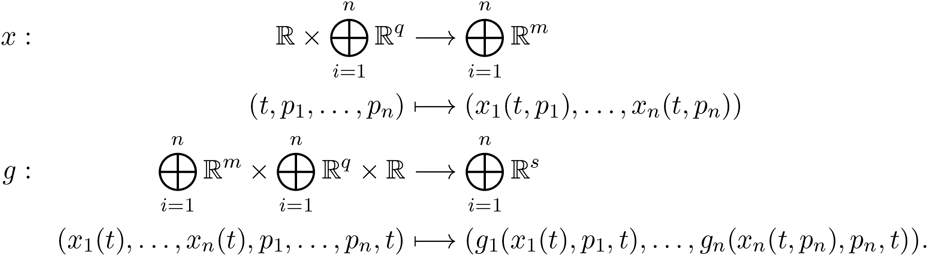

In words, if prediction- or observation functions are defined for different conditions then they expect condition-specific inputs which are evaluated by the matching functions. Examples for condition-specific prediction functions typically involve different forcings or events. Observation functions can e.g. differ between different measurement techniques. In all these cases, the different prediction functions *x*_1_, …, *x_n_* or observation functions *g*_1_, …, *g_n_* are defined, referencing the condition, and combined by the “+” operator analogously to `p`.

All commands, `P(), Xs(), Y()`, etc. can be executed with `condition = NULL`. In that case, the corresponding returned function is generic and, if called for different conditions, the same identical function is evaluated with the condition-specific input. The other way round, if a function, say, `x` is defined for several conditions, its prediction can be evaluated only for a subset of conditions by `x(times, pars, conditions = myconditions)`.

### 3.6. The data structure

In **dMod** different experimental conditions are handled by lists. Parameter transformations, prediction- and observation functions stacked by the “+” operator return list objects. On the other hand, data.frames as they are used for linear modeling, mixed-effects modeling or plotting with **ggplot2** are highly convenient to organize the data. The class datalist provides the interface between **dMod**’s list structures and `data.frame` objects. A `datalist` is a list of data frames with identical structure: observable names, time points, measured values and measurement uncertainty.

Objects of class `datalist` are usually generated by the `as.datalist()` command from a `data.frame`. The factor variables in the data frame to be used as generators for the unique condition names can be passed by the `split.by` argument. The resulting list of data frames has an additional attribute `“condition.grid”`, a translation table between the condition names and the original factor variables which can be used for specification of parameter transformations or augmentation of predictions by descriptive columns.

### 3.7. Objective function

The aim of **dMod** is parameter estimation. The objective function to be minimized for this purpose is the weighted least squares function which is produced by the `command normL2(data, prdfn)` where `data` is a `datalist` object and `prdfn` is a prediction function. The objective function is the final link connecting the chain of parameter transformations, prediction- and observation functions to observations. It collects all derivative information and besides the objective value it also computes gradient and Hessian. The standard optimizer employed within **dMod** is the trust() optimizer from the **trust** package.

Objective functions can be added by the “+” operator, meaning the objective values, gradients and Hessians are accumulated by summation. Besides the standard function `normL2()`, **dMod** provides several other functions returning objective functions, e.g. `constraintL2()` or `datapointL2()`. They allow to define parameter priors or treat data points as parameters, as shown in Section 4. Thus, a typical objective function used for parameter estimation could be

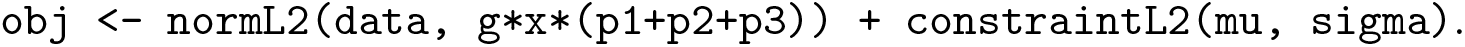

## 4. Three-compartment model of bile acid transport

The following model is a simplified dynamic model on bile acid transport. Bile acids are produced in the liver. They are necessary for the digestion of fat and oil. In the liver, bile acids are taken up in hepatocytes (liver cells) by specific transporter molecules. Clearance occurs either via canalicular or sinusoidal export, i.e. export to the bile transportation system of bile canaliculi or the intercellular space.

To study bile clearance, experiments with hepatocytes or hepatocyte-derived cell lines are performed *in-vitro*. Cells in a Petri dish stick together to a monolayer of cells, forming bile canaliculi in between cells. They adhere to the dish and are surrounded by a buffer providing the cells with nutrients. A radioactive label allows to measure the bile acid *taurocholic acide* (TCA).

For the mathematical description of bile flow, a three-compartment differential equation model is used. TCA is pipetted into the buffer compartment (`TCA_buffer`) from where it is transported into the cells (`TCA_cell`). Intracellular TCA is exported back to the buffer or the canalicular compartment (`TCA_cana`). Finally, canalicular TCA flows back into the buffer compartment. These processes give rise to the differential equations and corresponding flowchart presented in Figure 1.

**Figure 1:**
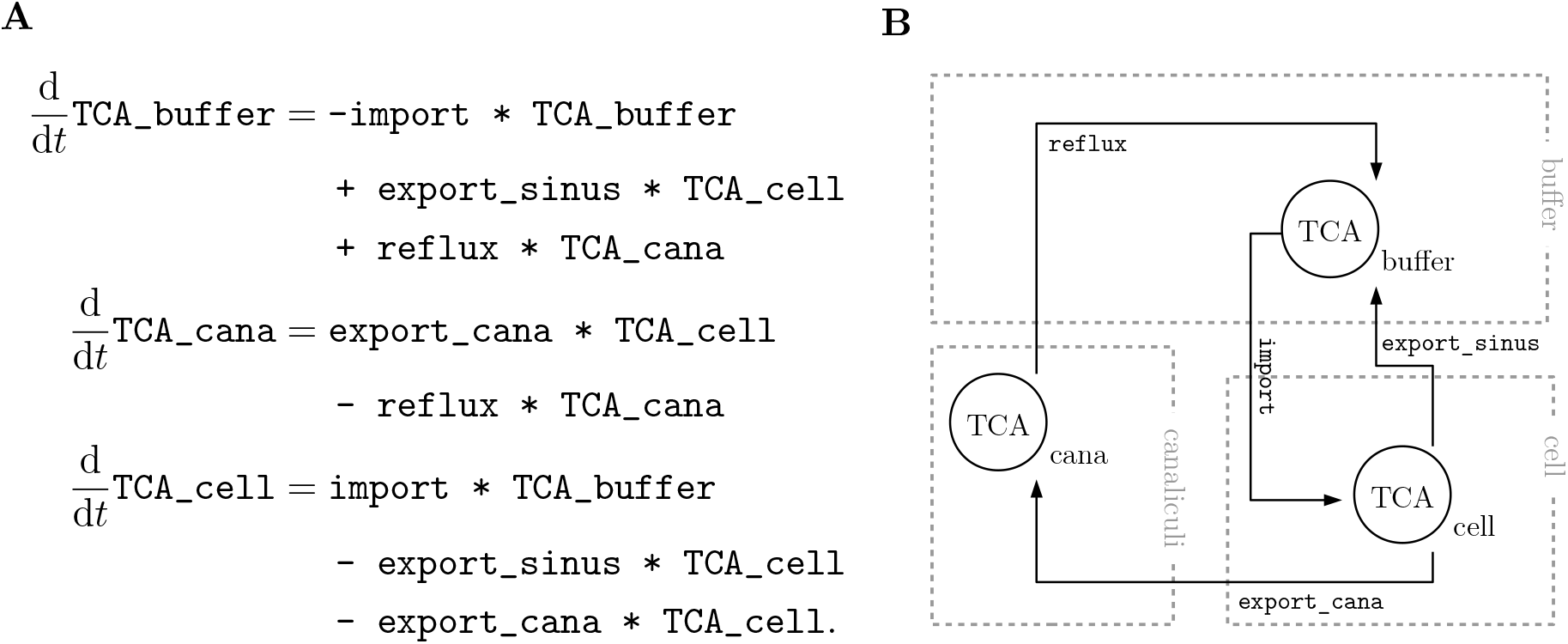
Differential equations and flowchart of the reaction network. Taurocholic acid, TCA, is transported between three compartments by four different processes. (A) Assuming mass-action kinetics, the three dynamic states satisfy a set of coupled differential equations. (B) The equations are visualized in a flowchart.

### 4.1 Simulation and prediction

Each transportation process is modeled by mass-action kinetics. A possible implementation in **dMod** is:

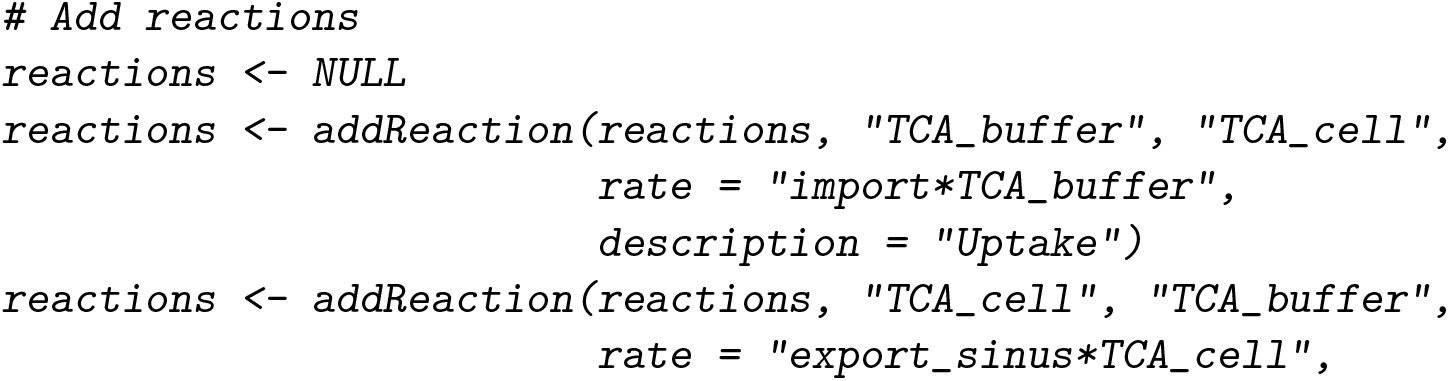

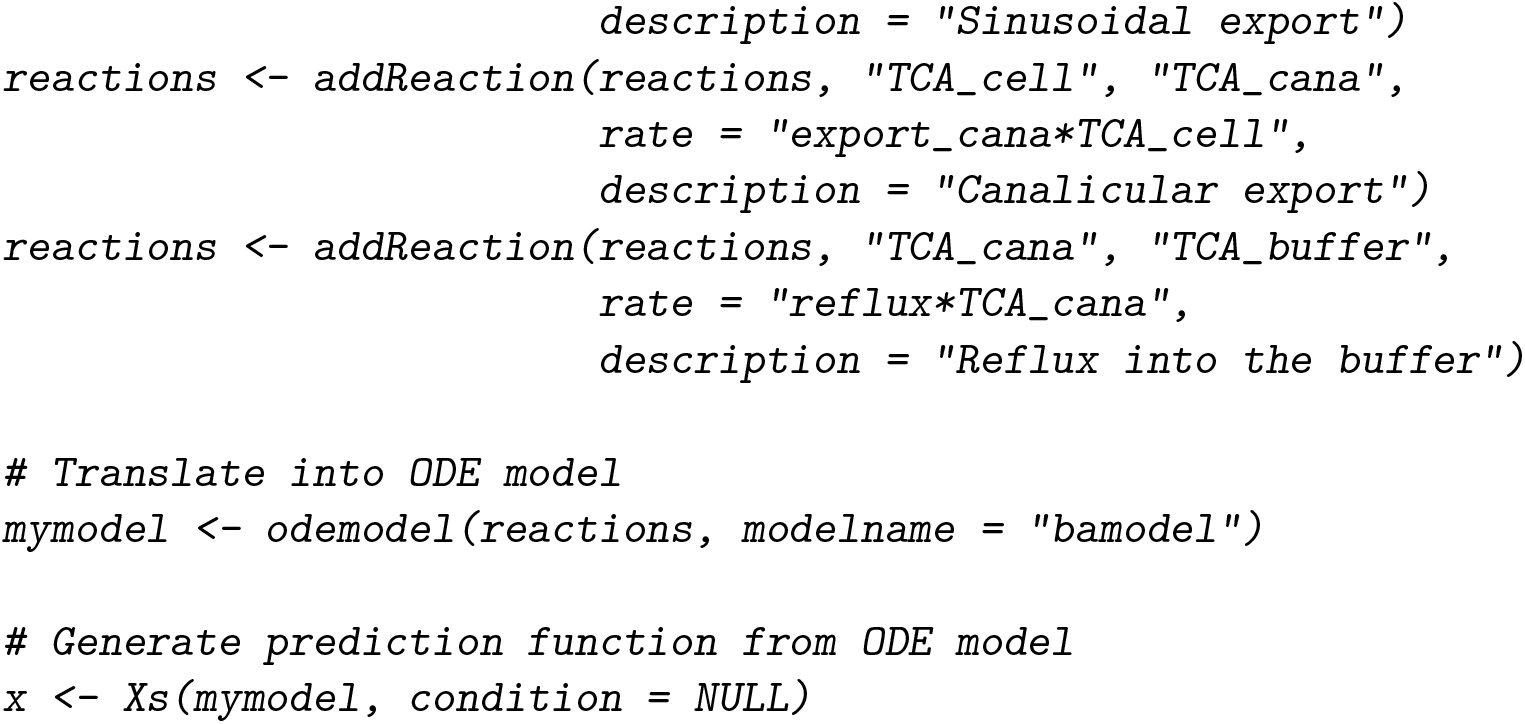

The reactions are collected in an `eqnlist` object. The `odemodel()` command composes the single reactions to an ODE system and auto-generates the C code which is used by the **deSolve** package to evaluate the ODE. Prediction functions are generated by the `Xs()` command. The usage of the prediction function is illustrated by the following code chunk. Time points are defined between 0 and 50, numeric values are assigned to all model parameters. Finally the prediction function is called and both, the prediction and the sensitivities, are plotted, shown in Figure 2.

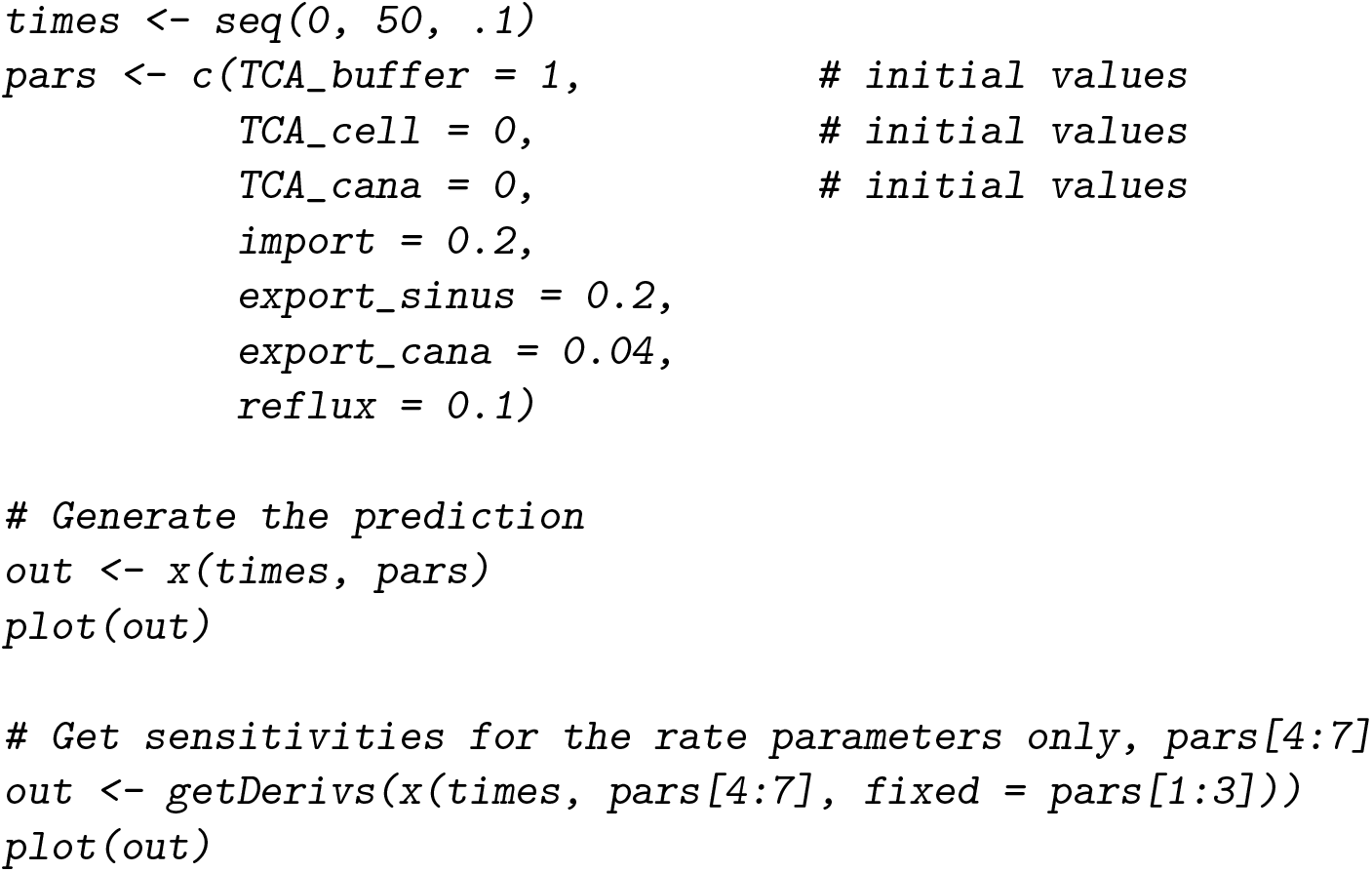

Figure 2A shows the uptake of `TCA_buffer` in the cell and canaliculi, saturating around *t* = 50. The prediction parametrically depends on initial values and rate parameters. Figure 2B shows the model sensitivities 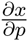 for the rate parameters.

**Figure 2:**
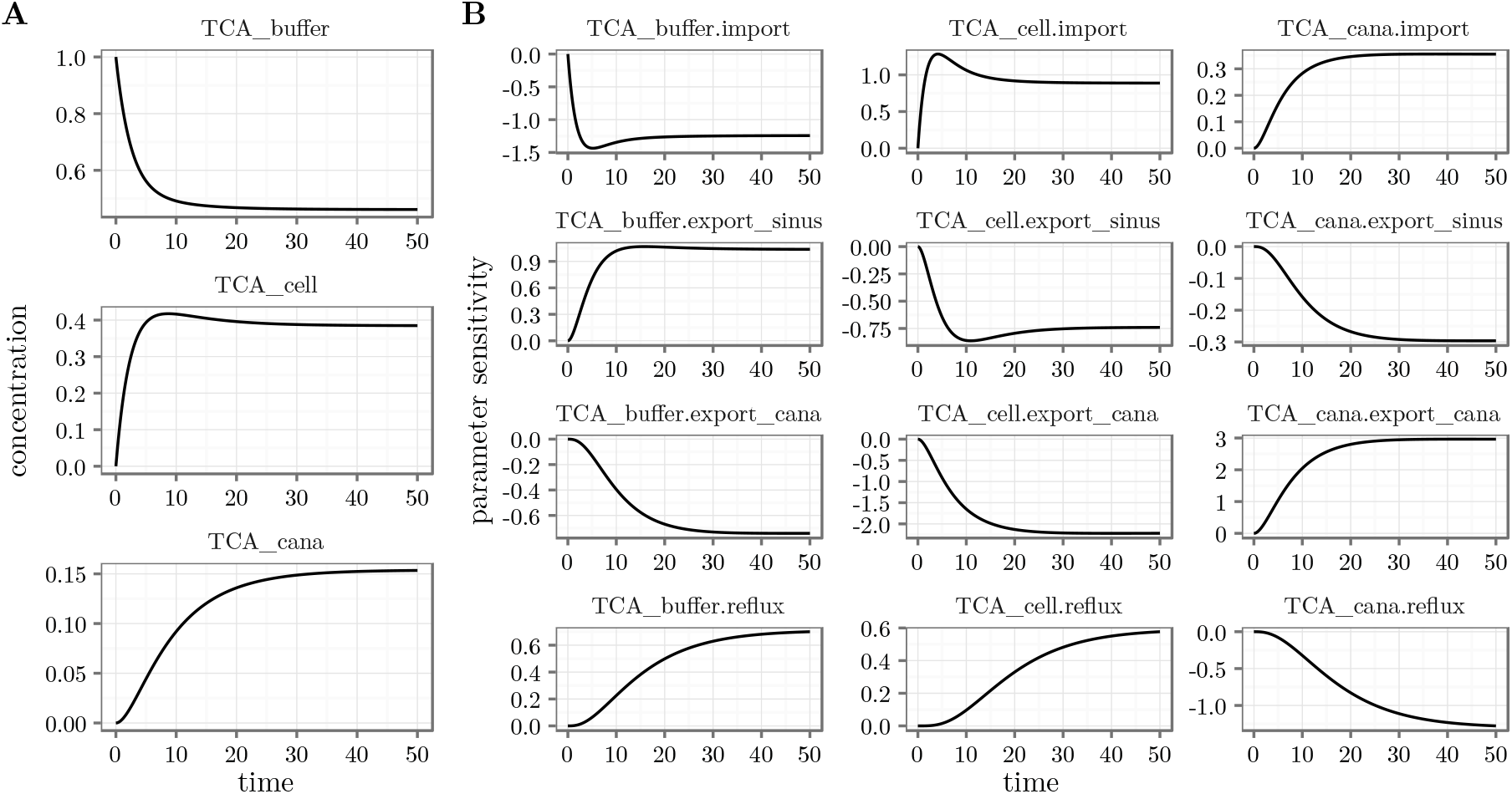
Output of the prediction function. (A) Prediction of the TCA states. (B) Sensitivities of the three TCA states for the rate parameters only.

### 4.2 Observation function and simulated data

In experiments, the three dynamic states, `TCA_buffer, TCA_cell` and `TCA_cana` cannot be directly measured. Rather, the radioactivity can only be measured separately for two compartments, namely the buffer and the cellular compartment, where the latter contains cells and canaliculi. This translates into the following relation between the radioactive counts and the dynamic states of our ODE model:

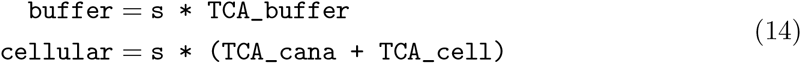

The scaling factor s translates amounts of TCA into radioactive counts. The observation function is expressed in **dMod** as follows.

First, observables are defined by an `eqnvec` object from which an observation function `g` is generated by the `Y()` command. The `Y()` command needs to be informed which of the symbols are variables (dynamic states) or parameters. Conveniently, `Y()` can parse an `eqnvec` or `eqnlist` such as `reactions` to retrieve this information.

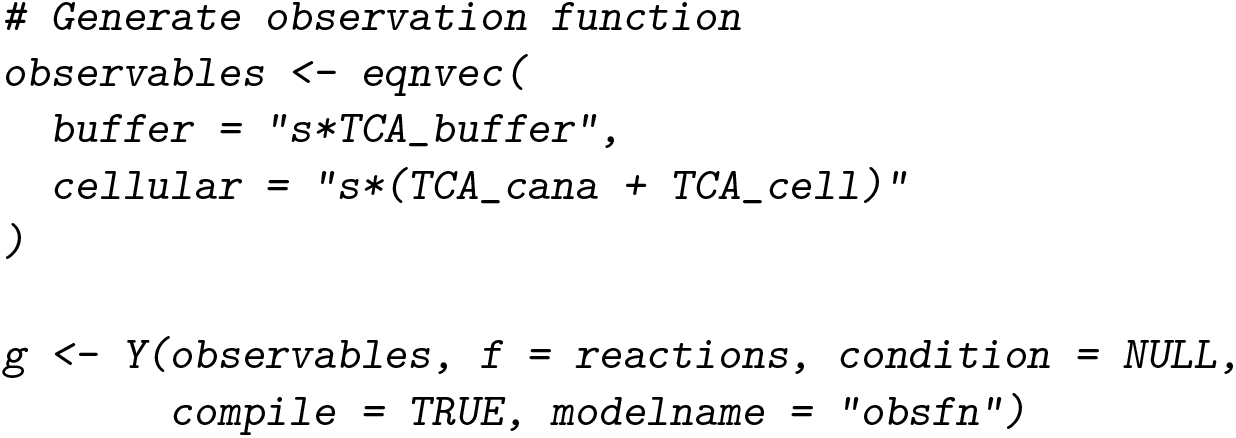

Observation functions link internal to observable states. Thus, providing values for the model parameters, the observation function can be used to simulate the outcome of an experiment. Adding noise to the prediction, experimental data is simulated. In the following, we will simulate the outcome of an *efflux experiment*. The experiment start with all TCA concentrations in steady state, such as shown in Figure 2 after *t* = 50. To initiate the efflux, the buffer is replaced by TCA-free buffer, i.e. `TCA_buffer = 0`. This translates into the following initial parameter values:

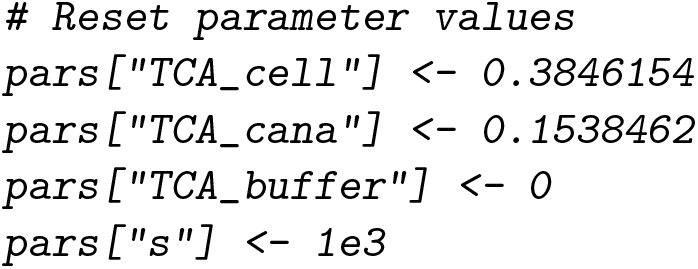

The predicted dynamics of the system’s internal and observable states is obtained by evaluation of the concatenated prediction function 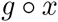, formulated as g*x in **dMod**. The scaling parameter s is set to 1000.

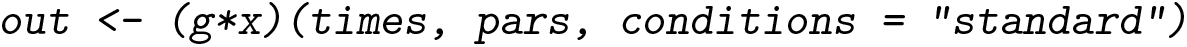

Since g and x have been generated as generic functions, i.e. `condition = NULL`, we can assign the output to a condition of our choice, in thas case “standard”. The predicted noiseless observation is obtained by considering the observable states only at the time point of observation timesD.

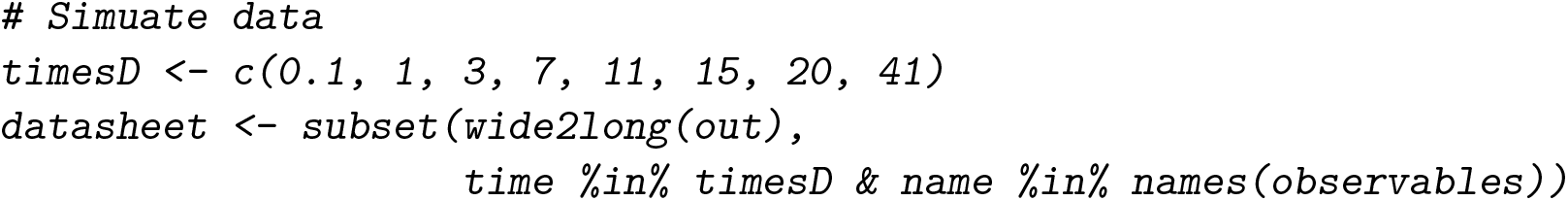

Data uncertainties *σ*are derived by the Poisson nature of radioactive count experiments, i.e. 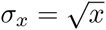. To avoid division by 0, the minimal *σ*-value is set to 1. Random values are added to the predicted values to simulate observation noise. In the end, the `data.frame` is converted into a `datalist` object.

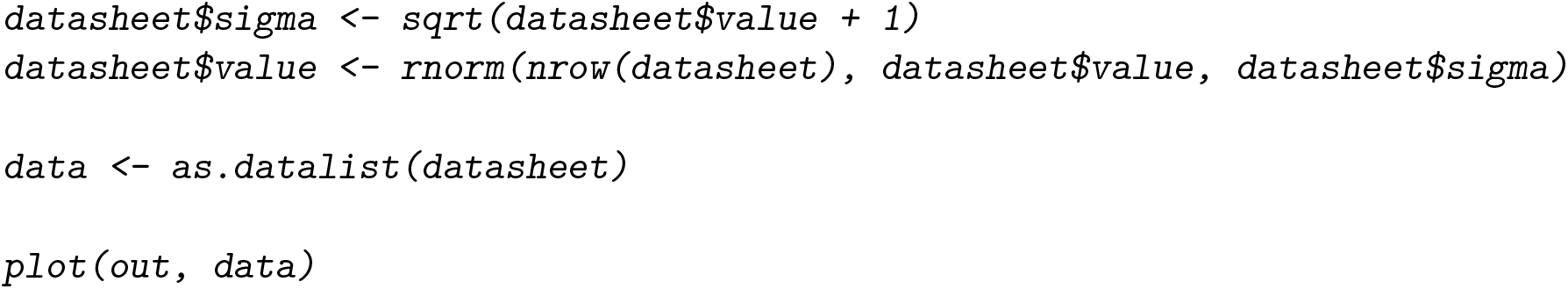

Both, the simulated data and the model prediction from which the data is derived are shown in Figure 3. The data reflects a typical time course of an efflux experiment, showing decreasing cellular TCA levels and increasing levels of TCA in the buffer.

**Figure 3:**
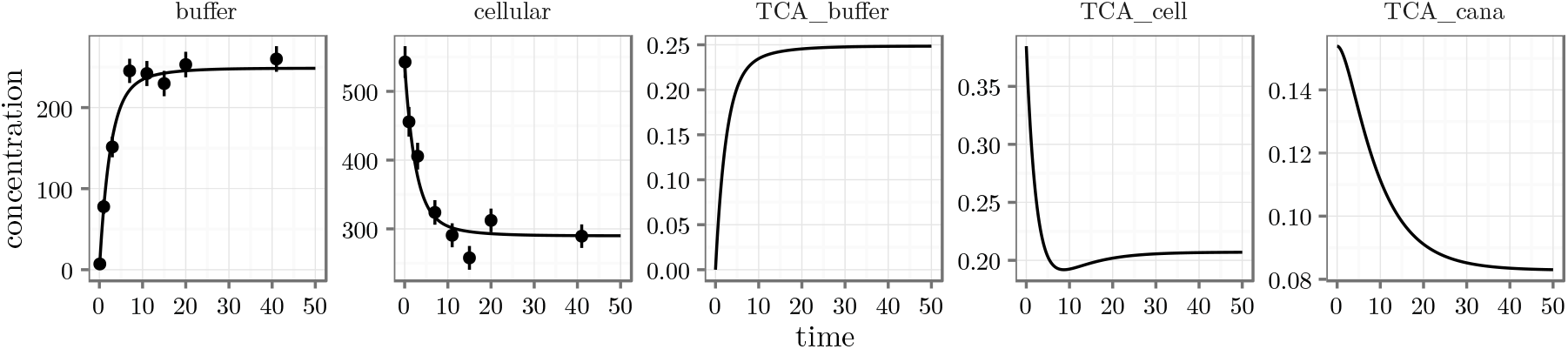
Model prediction of the observable and internal states. Simulated data is shown as dots with error bars.

### 4.3. Parameter transformation

Parameter transformations play a crucial role in the set-up of **dMod**. They can have several purposes such as fixing paramer values, implementing parameter bounds, including steady-state constraints or mapping parameters to different conditions. While being conceptually the same, it might be worth noting that parameters can be distinguished in two classes. The first class of parameters are *initial values* for dynamic states, such as `TCA_buffer`. Parameters of the second class, such as rate parameters, have no accompanying dynamic state.

First, we use parameter transformations to constrain all parameters to be positive or zero because all our parameters are either amounts or rate parameters. The parameter transformation is generated by the `P()` command taking an `eqnvec` object. Parameter transformations explicitly state the relation between the **inner** parameters, i.e. the parameter values that are evaluated within the model, and the **outer** parameters, i.e. the parameter values provided by the user or by an optimizer. In our case, we imply positivity of inner parameters using the `exp()` function on outer parameters. The corresponding code reads:

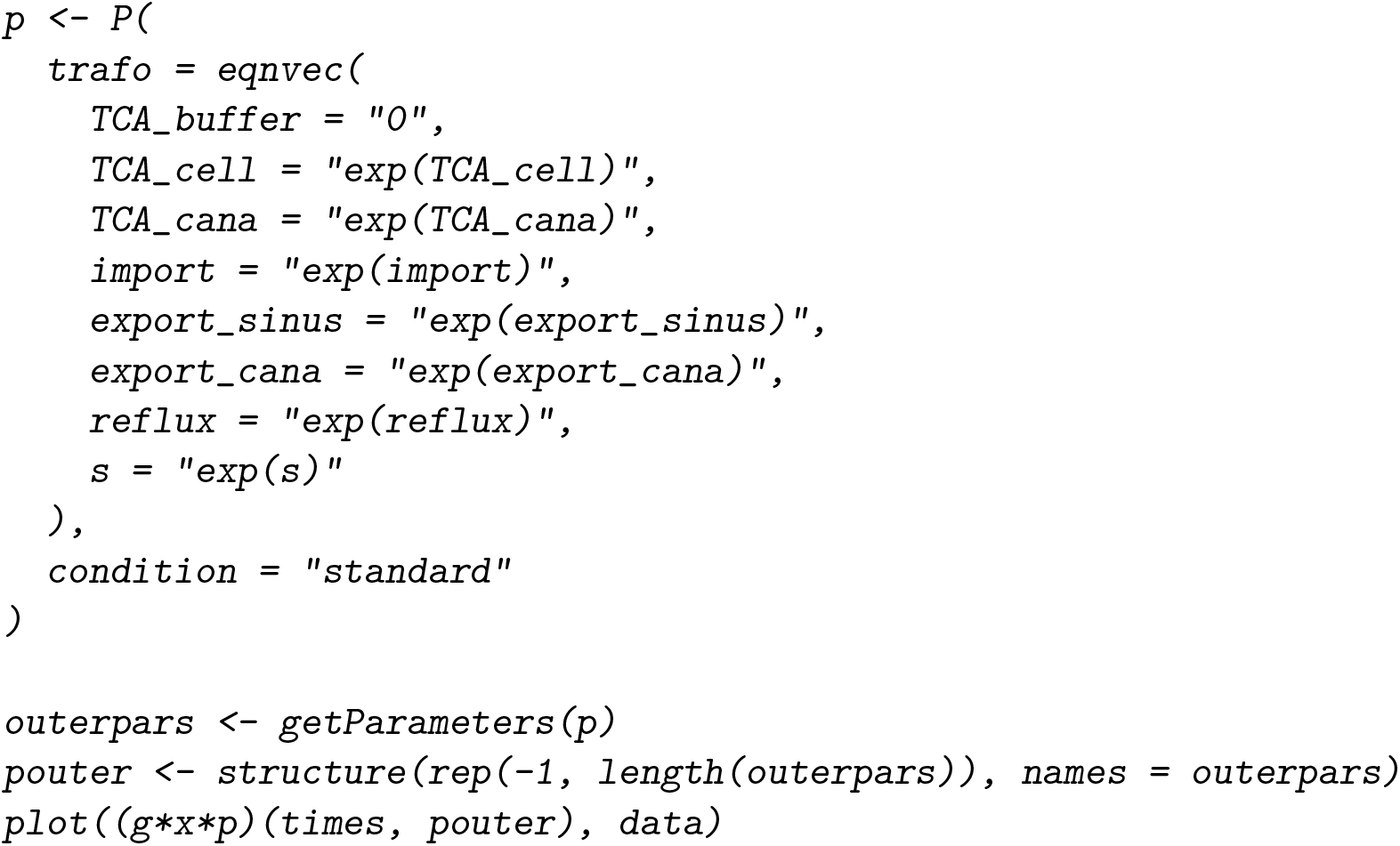

The vector `outerpars` is the collection of all symbols on the right-hand side of trafo. It coincides with `names(pars)` except for `TCA_buffer`, which is fixed to a constant expression, here 0, by the transformation. However, the interpretation of the parameters has changed since now their values are on a log-scale. All three functions, the observation function, prediction function and parameter transformation can be concatenated to one new prediction function, g*x*p which takes times and values of the outer parameters to predict internal and observable states. The model prediction generated by pouter is shown in Figure 4.

**Figure 4:**
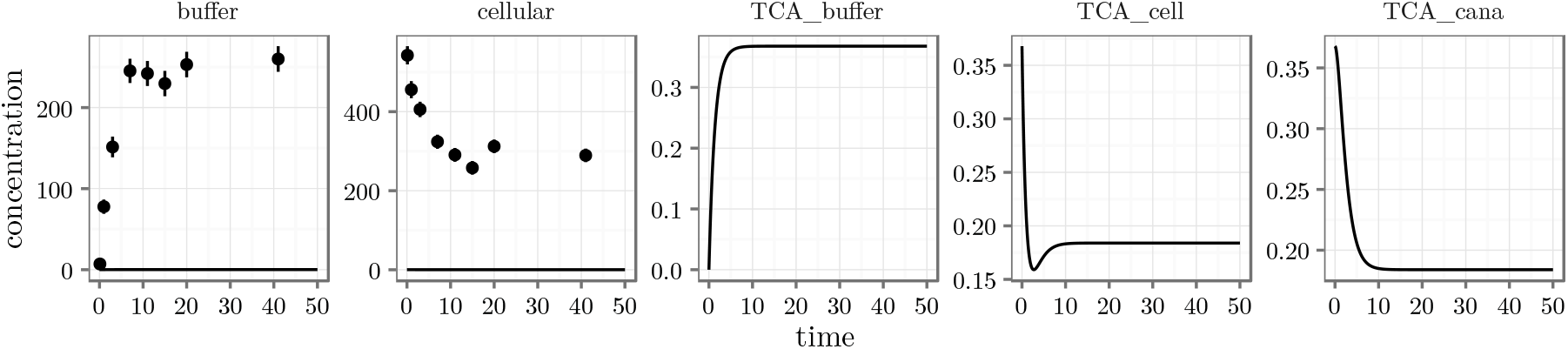
Prediction of internal and observed states. All values of the outer parameters have been set to –1. Simulated data points are shown as dots with error bars.

### 4.4. Objective function and model fitting

The objective of model fitting is to find parameter values such that the corresponding model prediction matches the observation. In Figure 4, the graphs of `buffer` and `cellular` should match the observation within the error. For normally distributed measurement noise, maximum-likelihood estimation is equivalent to least-squares estimation. A least-squares objective function can be generated by the `normL2()` command which requires a `datalist` object, in our case `data`, and a prediction function, in our case `g*x*p`.

Frequently, non-identifiable parameters are encountered in non-linear dynamic systems. To prevent the optimizer from selecting extreme parameter values for which the ODE solver aborts, a general quadratic prior can be entailed on all parameters by calling `constraintL2()`. The objective functions returned by `normL2()` and `constraintL2()` are objects of class objfn and can be added by the “+” operator.

The following code illustrates the implementation of the objective function and how it is used with the `trust()` optimizer from the **trust** package to obtain a model fit, shown in Figure 5.

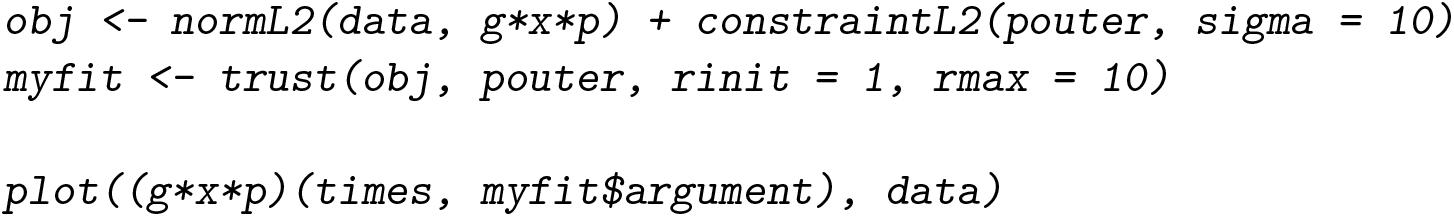

Besides non-identifiability of parameters, local optima constitute another pit-fall when optimizing non-linear functions. `The trust()` optimizer employs derivative information and therefore, if starting within a certain region around a local optimum, is very efficient in finding it back. Once an optimum is found, we can be confident that there is no deeper point around. However, to be confident that an optimum is the globally best solution, we might want to scatter starting points for optimization runs all over the parameter space. The **dMod** package provides the `mstrust()` function based on `trust()` to do a multi-start search:

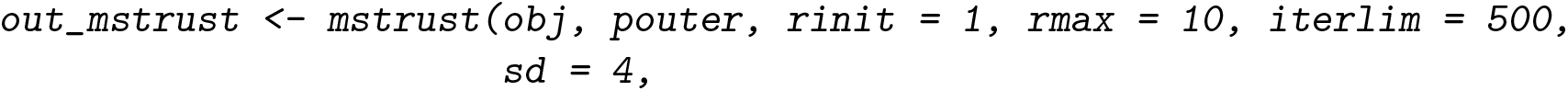

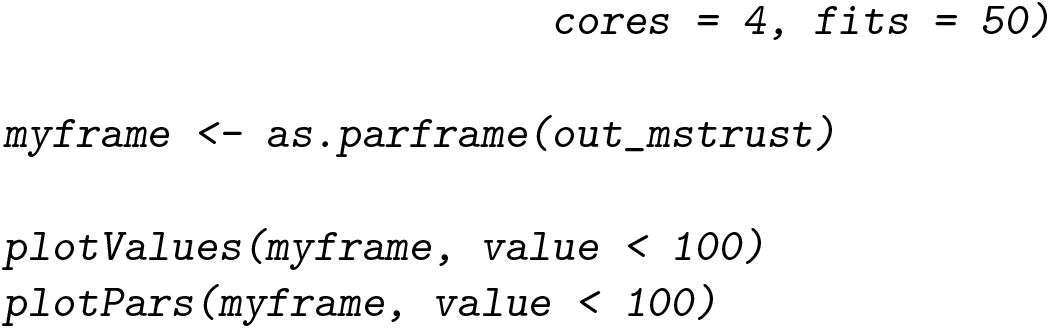

**Figure 5:**
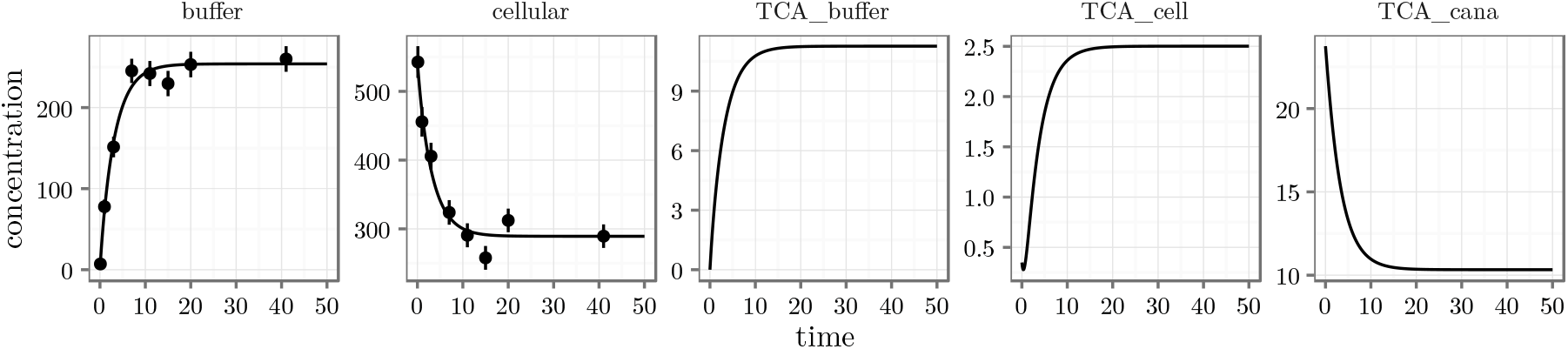
Prediction of internal and observed states after optimization of the objective function. Simulated data points are shown as dots with error bars.

Here, we have searched according to 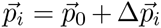 is the center, in our case pouter, 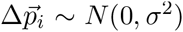 is a random parameter vector taken from a normal distribution, in our case *σ* = 4, and the index *i* runs from 1 to fits = 50. The `mclapply()` command from the **parallel** package is used internally to run fits in parallel, here `cores = 4`. The result of `mstrust()` is a list of all returned values of `trust()`. To extract the final objective value, parameter values, convergence information and the number of iterations, `as.parframe()` is used. The multi-start approach identifies four local optima, see Figure 6, which yield almost the same objective value, Figure 6A. Despite the similar objective value, the optima are not close to each other in parameter space, as being illustrated by Figure 6B, and lead to different predictions, Figure 6C. Figure 6B suggests that the two initial values `TCA_cell` and `TCA_cana` are connected in the sense that if one takes a large value, the other takes a small value and vice versa. This is not surprising because the observed cellular TCA amount is the sum of both. A new experiment needs to be designed to distinguish one situation from another.

**Figure 6:**
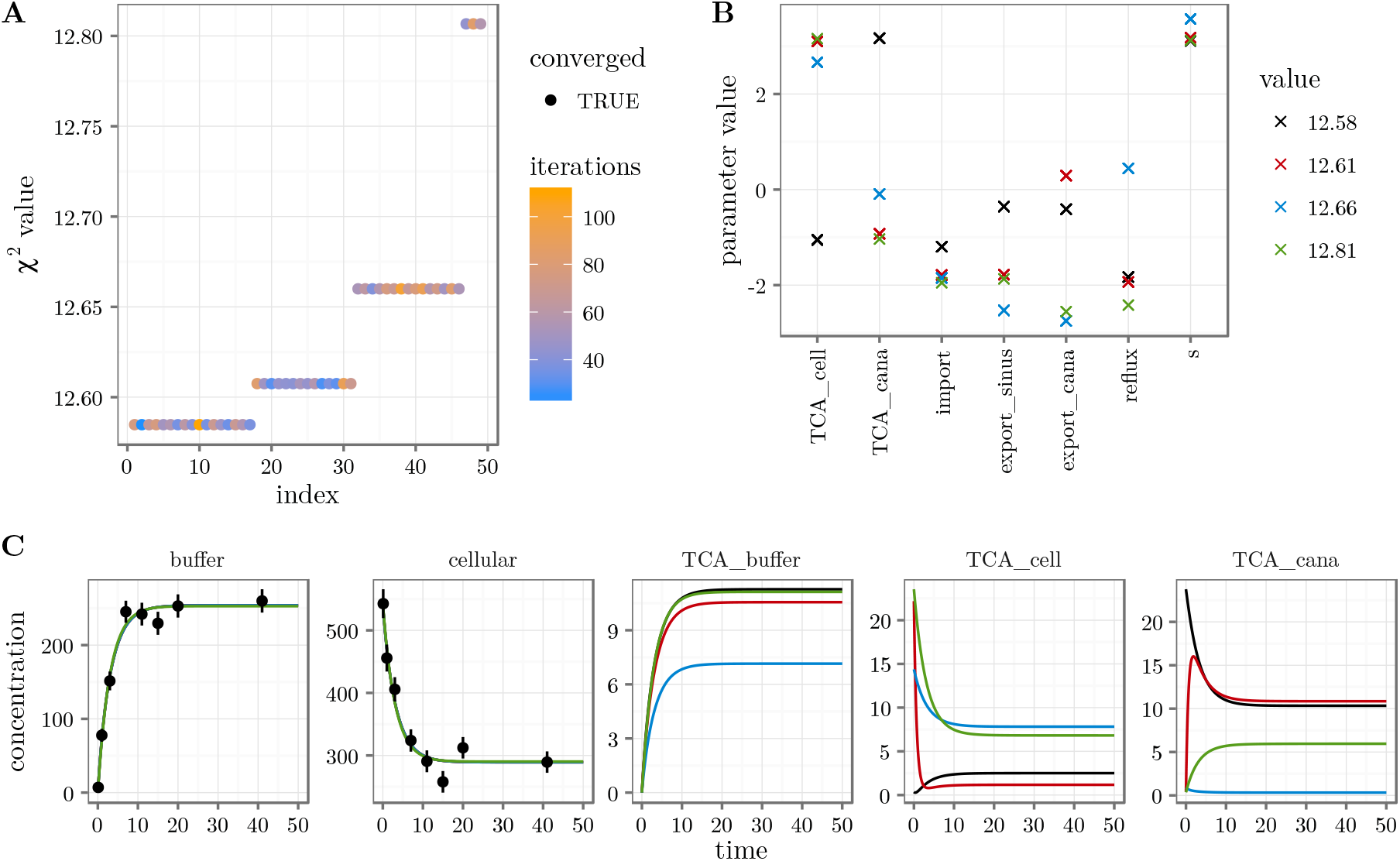
Result of multi-start fitting procedure. (A) Fits have been sorted by increasing objective value. Four optima were found with almost identical objective value. (B) The parameter values for different optima are shown in different colors. (C) Each local optimum corresponds to a different model prediction, shown in different colors. The observed states are pracitically undistinguishable although the internal states show different behavior.

### 4.5. Working with several conditions

In practice, the canaliculi only form a closed compartment if Ca^2+^/Mg^2+^ ions are present in the buffer. Therefore, if the experiment is repeated with Ca^2+^/Mg^2+^-free efflux buffer, the contents of the canaliculi escapes quickly into the buffer compartment. Under this condition, the `buffer` measurement reflects what was formerly the total TCA content in buffer and canaliculi whereas the `cellular` measurement reflects what was fomerly the TCA content of the cells. Mathematically, two experimental conditions which differ only by the reflux parameter need to be combined in one objective function.

Like before, we simulate a data set. Then, a parameter transformation for the additional condition is set up and the parameter space is explored by a multi-start fit.

To simulate the new experimental condition, the ‘reflux’ parameter is modified. The new data set is combined with the original data by the “+” operator.

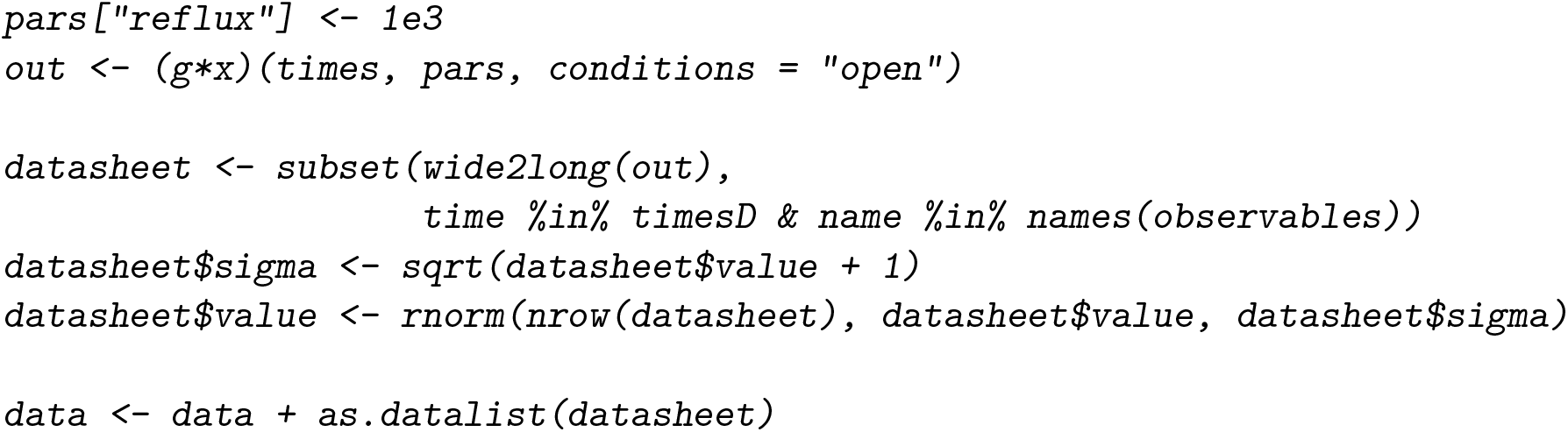

To add a condition to the parameter transformation function, we use the equations of the standard condition as a template for the “open” condition. Parameter transformation functions for different conditions are combined by the “+” operator.

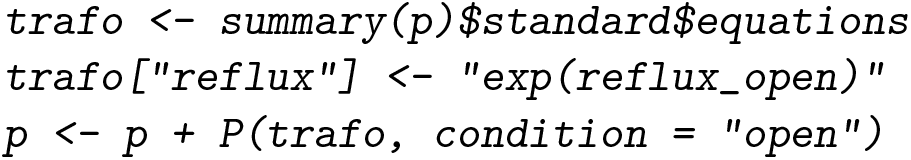

Both transformations “standard” and “open” now possess the outer parameters `reflux` and `reflux_open`. However, the value of `reflux` is mapped to an inner parameter only by transformation “standard”. Accordingly, transformation “open” only uses `reflux_open`. Thus, both transformations return the same values for all but the `reflux` parameter. The prediction function g*x is generic in the sense that `condition = NULL` whereas the concatenation `g*x*p` has the conditions “standard” and “open”, evaluating the identical function g*x on two parameter vectors.

We define an updated objective function:

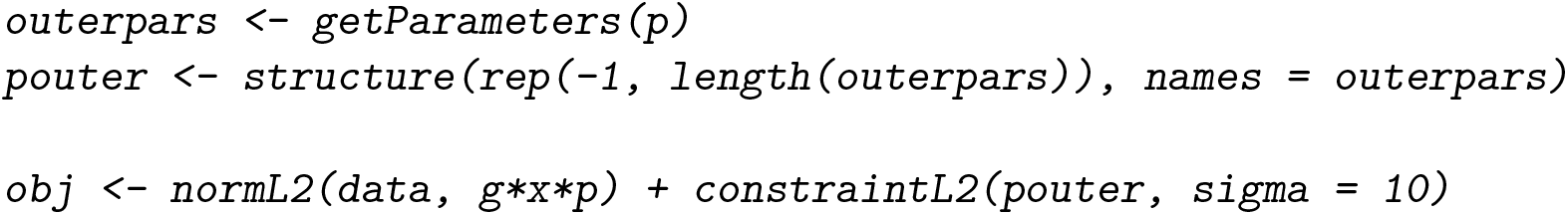

Then we start 50 fits around `pouter`. The list of fits is simplified to a `parframe` and by the `as.parvec()` function, the parameter vector (of the best fit) is extracted from the `parframe`. The best fit is used to make a prediction which is plotted together with the simulated data. All results are shown in Figure 7.

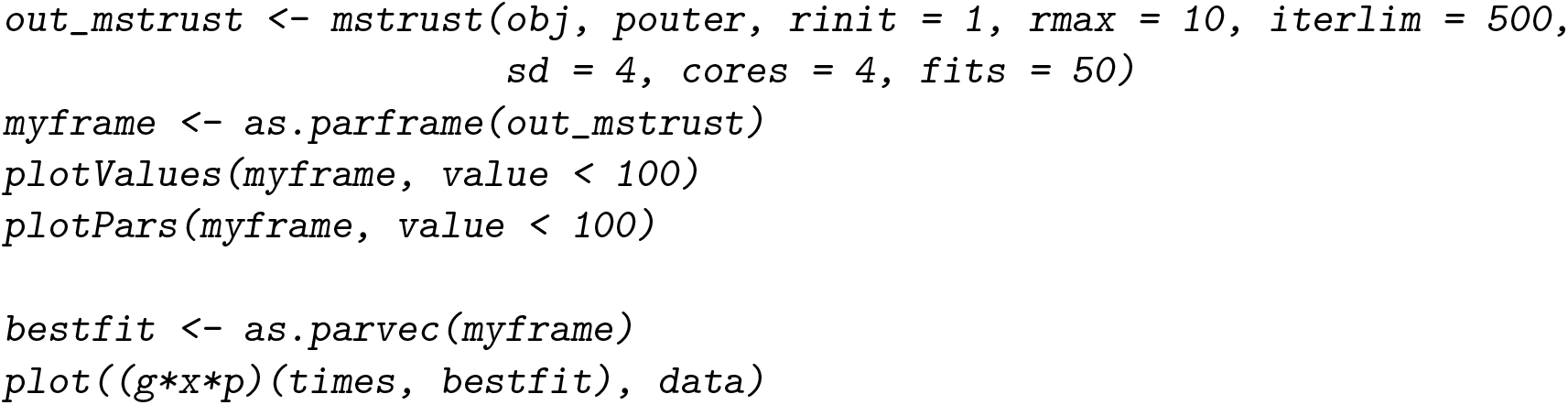

**Figure 7:**
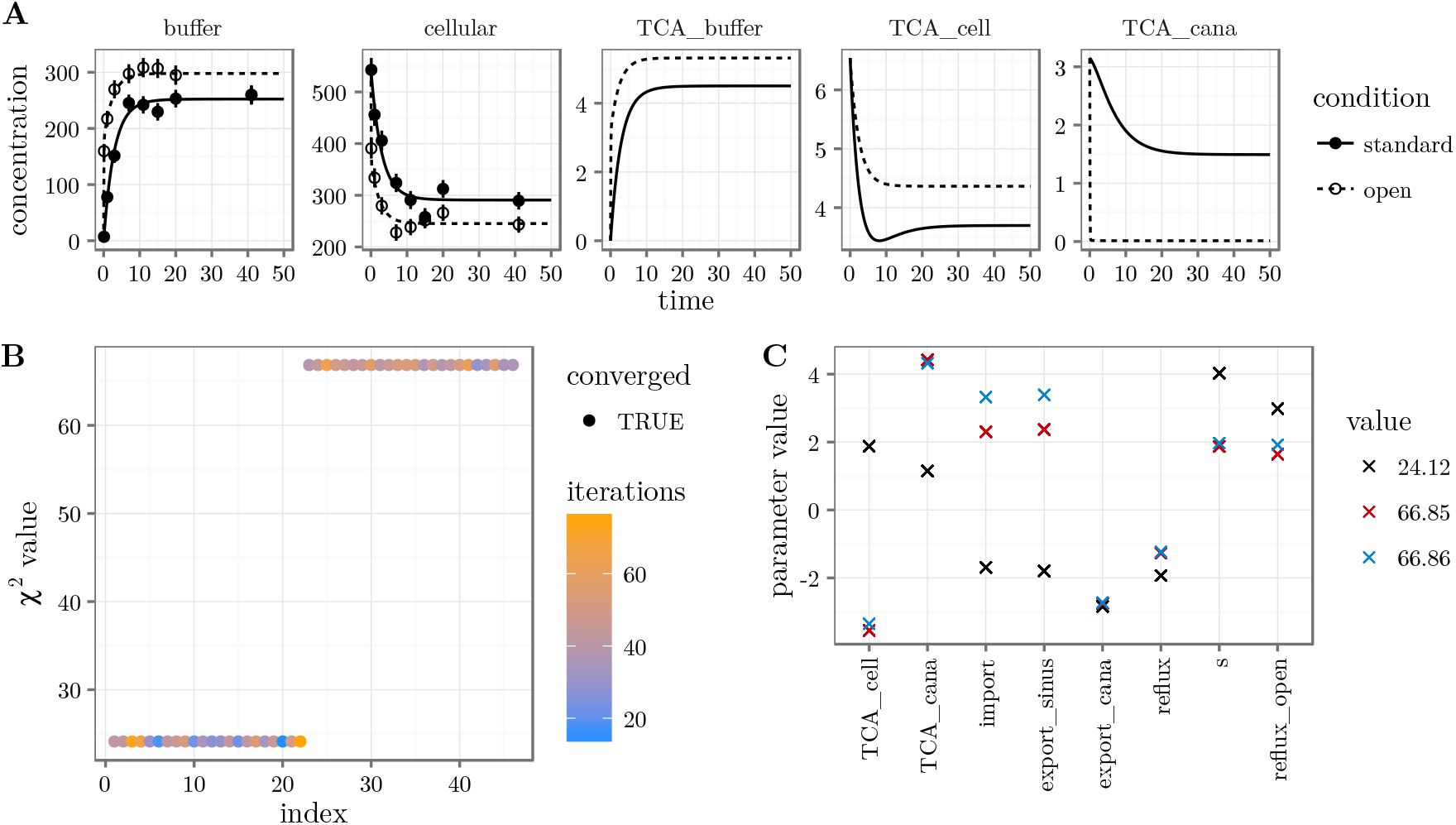
Result of multi-start fitting procedure with two experimental conditions. (A) Model prediction of the best fit and the simulated data are shown in different colors. (B) Fits have been sorted by increasing objective value. The lowest value clearly separates from the second plateau. (C) Plotting the parameter values for each of the fits reveals that the second plateau consists of two optima. The lowest plateau however corresponds to a unique optimum.

Interestingly, with the new experiment the best optimum becomes unique. The log-likelihood difference, i.e. half the difference between the objective values, is more than 20 between the lowest and the second plateau, which is highly significant. The uniqueness of the lowest plateau is confirmed by Figure 7C which shows no scattering of the black circles.

### 4.6. Parameter uncertainty and identifiability

One might wonder why the optimum is unique as for any choice of the scaling parameter s we find appropriate values of the TCA initial value parameters that give rise to exactly the same prediction of the observables. The reason for the uniqueness is the parameter *L*2- constraint that we have added to the objective function. Nonetheless, we will see, that the non-identifiability is still visible in the profile likelihood.

The profile likelihood is computed by the `profile()` command. There are several options to control the step-size and accuracy. For convenience the `method` option can be used to select between the presets `“integrate”` and `“optimize”`.

The code

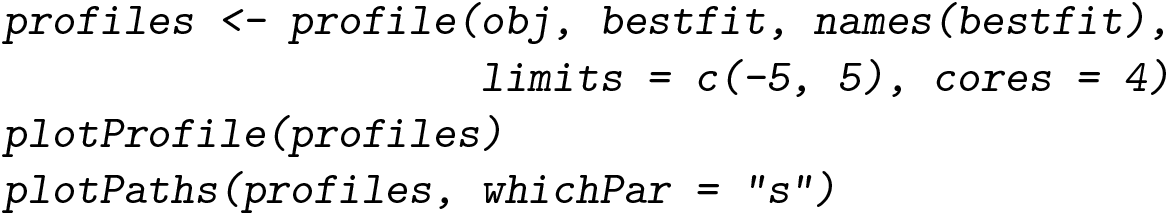

gives us the result shown in Figure 8. Computing the profile likelihood, the sum of data contribution, `normL2`, and prior contribution, `constraintL2`, are optimized under the constraint of a given parameter value for the profiled parameter. In the optimum, data and prior contribution are evaluated separately giving rise to the dashed and dotted lines in Figure 8A. As we had expected, the data contribution to the initial value parameters `TCA_cell, TCA_cana` and the scaling parameter s is constantly zero. The parameters are structurally non-identifiable.

**Figure 8:**
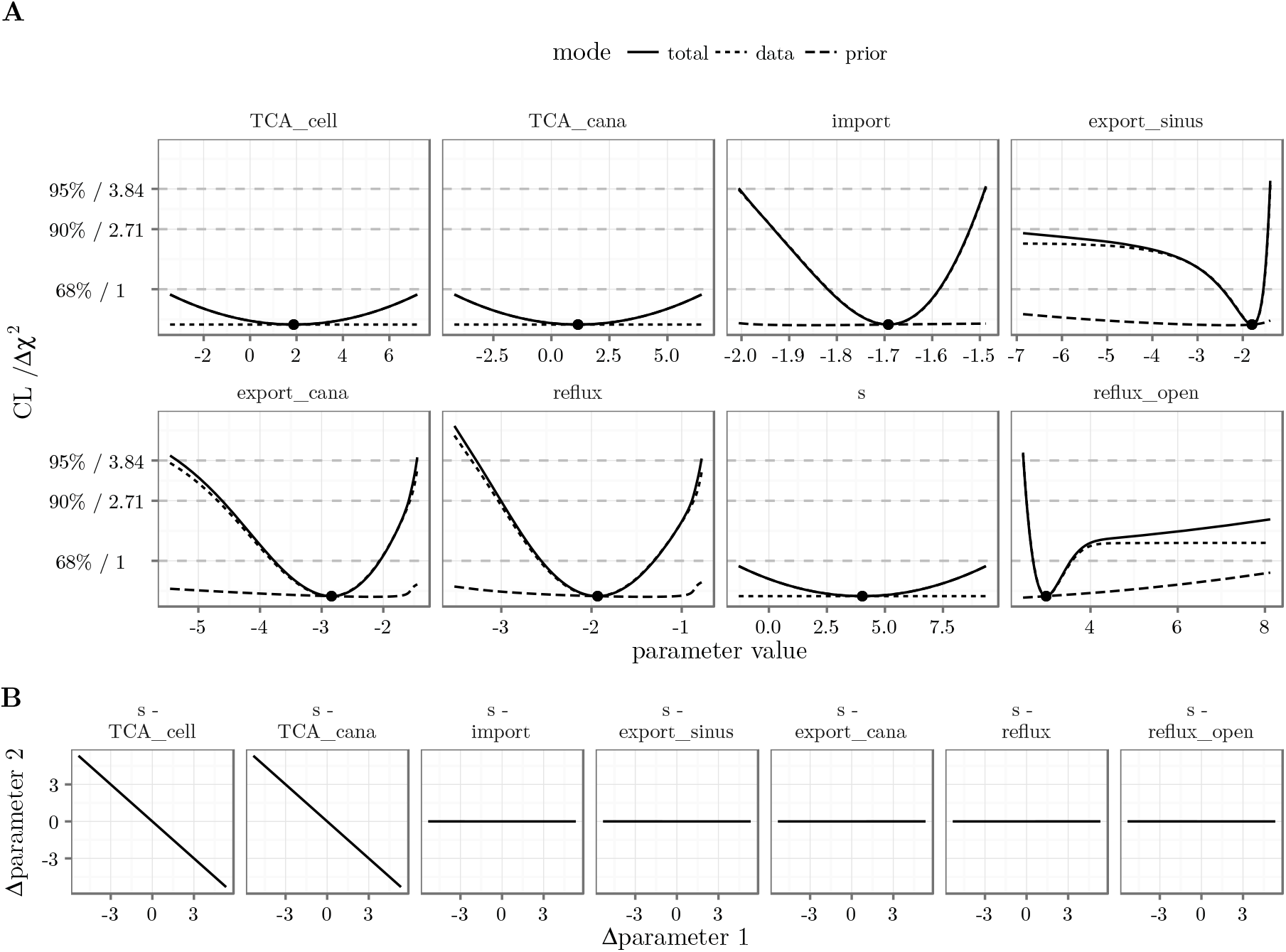
Profile likelihood. (A) Profiles of all parameters. Data- and prior contribution to the total objective value are distinguished by line-type. (B) Parameter paths for the scaling parameter s.

Each profile corresponds to a certain path in parameter space. The path for the profile of the non-identifiable scaling parameter s is shown in Figure 8B. It shows a clear coupling of the scaling parameter `s` and the initial value parameters `TCA_cell` and `TCA_cana`: both initial value parameters have to be decreased by the same extent as the scaling parameter is increased to keep the prediction unchanged.

The profiles of the parameters `export_sinus` and `reflux_open` exceed the 95% confidence threshold only to one side. Given the data, the `export_sinus` parameter could equally be –∞ (corresponding to an export rate of 0) without changing the likelihood significantly for the worse. A similar statement holds for the `reflux_open` parameter which could equally be ∞ meaning that we could assume instantaneous draining of the canaliculi for the `“open”` condition. The two parameters are practically non-identifiable.

Finally, the parameters `import, export_cana` and `reflux` exceed the 95% confidence threshold in both directions meaning that the parameters have finite confidence intervals. However, the confidence intervals are rather large and we might ask if there is further information that we could use to improve parameter identifiability without generating new data.

### 4.7. Steady-state constraints and implicit transformations

So far we have estimated both initial concentrations, `TCA_cell` and `TCA_cana`, independently. However, we know that the efflux experiment was just started after completion of the uptake process. Our system runs into a steady state, the buffer is exchanged and the measurement begins. Hence, we can use the steady-state condition as an additional information for the modeling process.

The steady-state relation between `TCA_cana` and `TCA_cell` can be derived analytically from the ODE, Figure 1A, by setting the right-hand side of 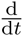 `TCA_cana` to zero. It reads

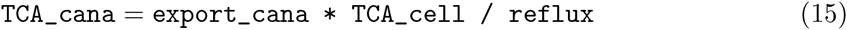

This relation can be explicitly used in a parameter transformation to express `TCA_cana` in terms of other parameters. The dimension of the parameter space is thereby reduced by one. The following implementation shows how we would use the existing transformation function p to generate an alternative transformation function pSS which includes the steady-state condition and replaces p in the prediction function g*x*p.

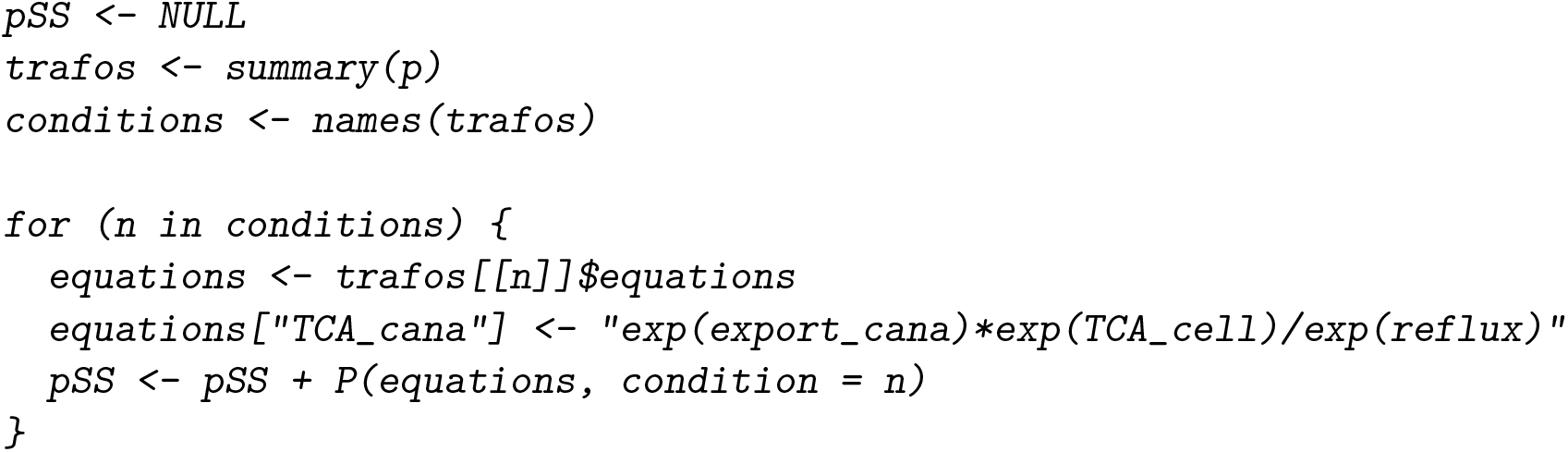

We get all the information about the transformations from the `summary()` command. The equation for `TCA_cana` is substituted by our steady-state constraint. By the “+” operator, a new parameter transformation function `pSS` is iteratively constructed for all conditions. Alternatively, we want to implement the steady-state constraint by an *implicit* parameter transformation, as opposed to the explicit transformation shown above. Let 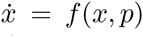 be our dynamic system. Then, under certain conditions, we find a function *g*(*p*) such that *f g*(*p*)*, p* = 0 for all *p*, i.e. *xS* = *g*(*p*) is a steady state of *f*. The parameter transformation we want to generate is the function 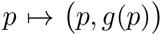. Here, the set of outer parameters is the set of the reaction rates *p* whereas the set of inner parameters contains these rates and the corresponding steady states as initial value parameters. The root of *f* must be determined numerically to which end `multiroot()` from the **rootSolve** package is used.

The following code is a reimplementation of the example above. The condition “open” is special in the sense that the reflux rate is modified compared to the reflux rate by which the steady state is computed. This is solved by an event, flipping a switch variable from 0 to 1 and thereby replacing the rate `reflux by reflux_open`.

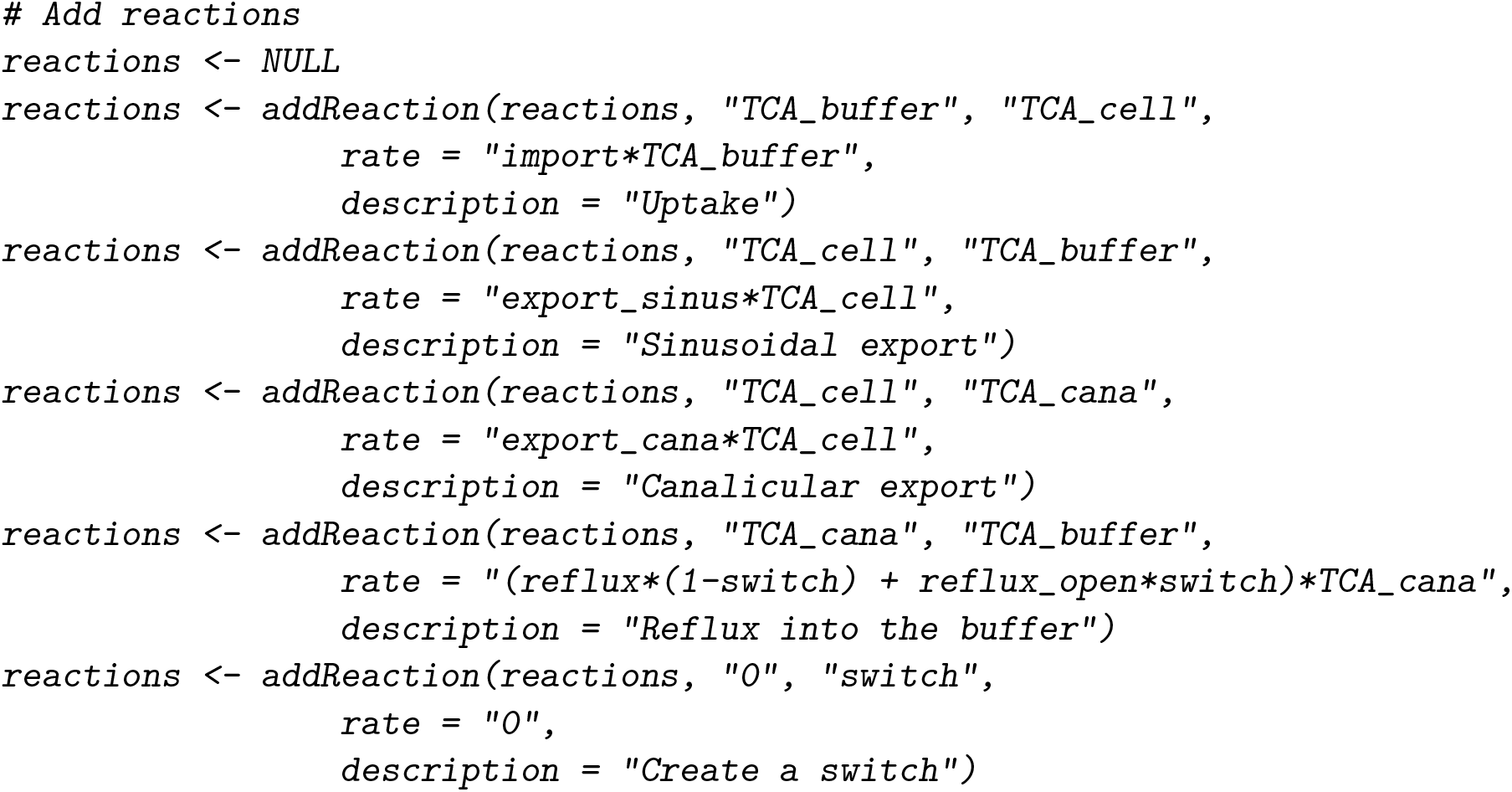

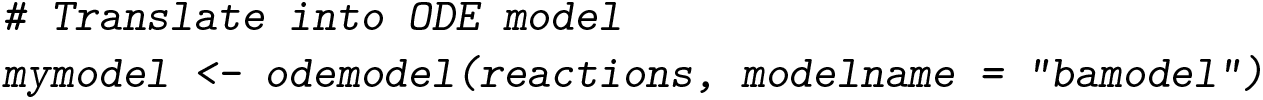

For the implicit parameter transformation we need the ODE which is obtained by the command `as.eqnvec()` from the reactions. The Jacobian of *f* is rank-deficient because the system has a conserved quantity *c* = TCA_buffer + TCA_cana + TCA_cell which is the total TCA amount. Replacing one element of *f* by *c* – `TCA_tot`, the rank of the Jacobian is completed, the condition for the local existence of the implicit function *g*(*p*) is satisfied and the steady state is parameterized by *p* and the additional parameter `TCA_tot`.

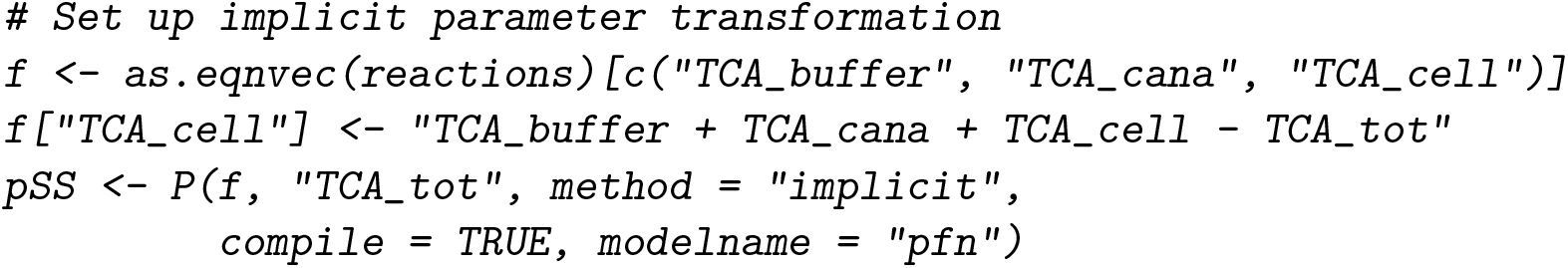

For the optimization, all outer parameters should still be log-parameters, implemented by an explicit parameter transformation. Outer parameters for the initial values are not necessary any more. They can be replaced by 0 since the initial values are computed by the implicit transformation. The final transformation will be a concatenation of the implicit and explicit transformations, pSS*p.

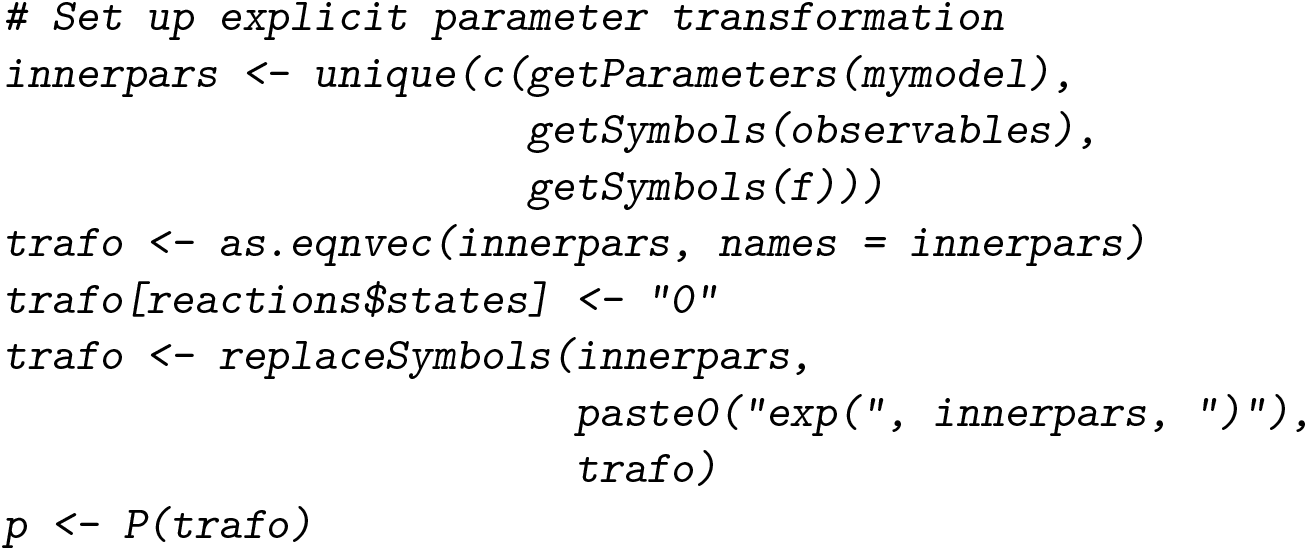

Both, the exchange of buffer and the opening of bile canaliculi by a Ca^2+^/Mg^2+^-free buffer are implemented as events. We therefore have different prediction functions for the two conditions.

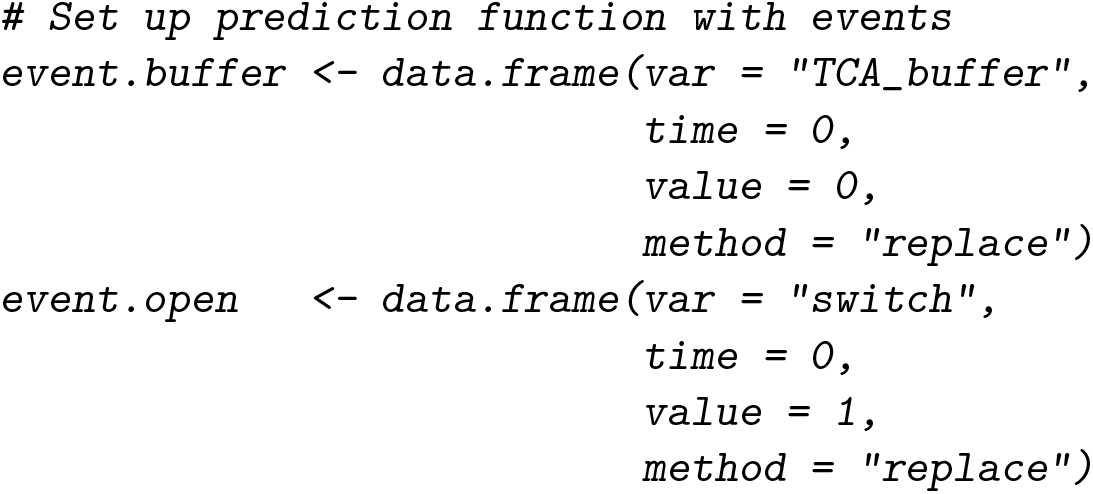

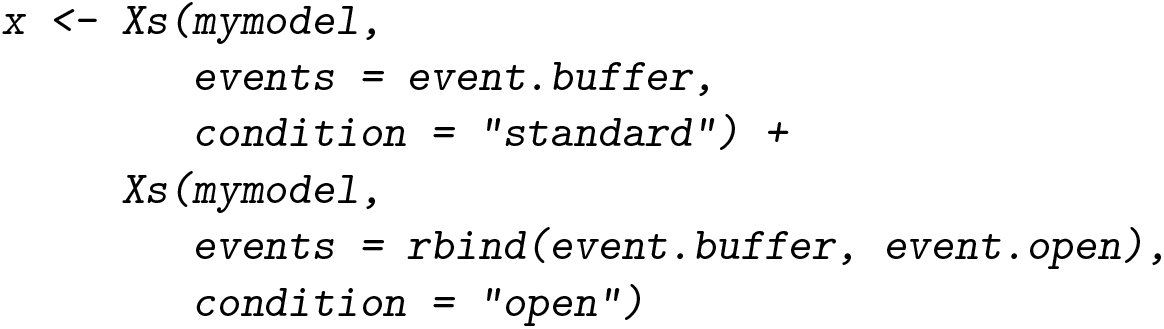

Although the observables have not changed compared to the set-up with purely explicit transformations, the observation function must be generated again because the new parameter `TCA_tot` has appeared. This parameter was not contained in the original reactions. Therefore, we use the states and parameters arguments to explicitly inform the observation function about the *states* and `parameters` involved. Otherwise, parameter sensitivities are not propagated correctly.

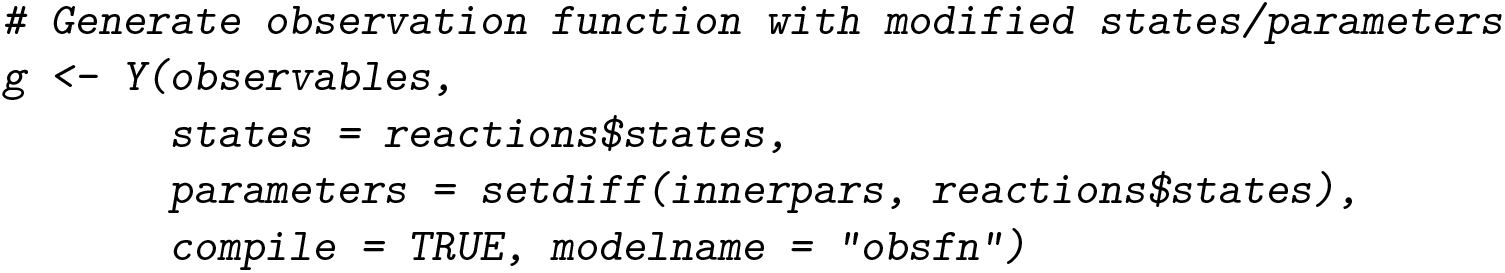

Finally, the objective function is defined. The prediction function is now a concatenation of four functions.

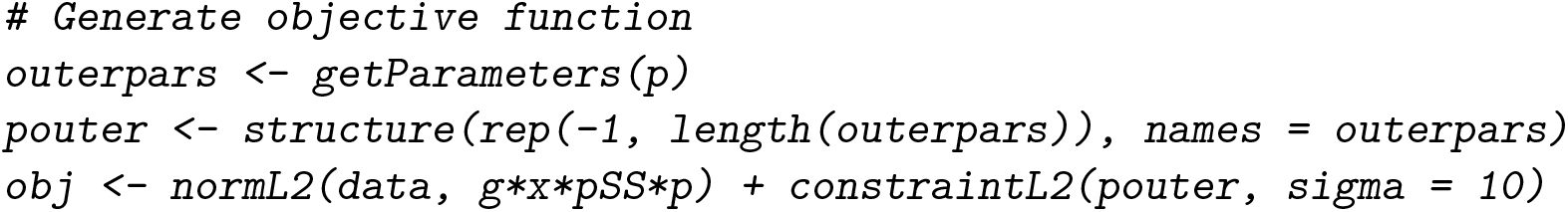

The same simulated data set has been fitted by the fully explicit and the implicit/explicit model implementations. In both cases the global optimum is unique. Parameter profiles have been computed with both model implementations, shown in Figure 9. In addition, the original profiles without steady-state constraints are plotted. The optima found by all three approaches, no steady-state, analytic steady-state and numeric steady-state, are statistically compatible. The two implementations using the steady-state information show exactly the same profiles. Since one formulation is parameterized by `TCA_cell` whereas the other is parameterized by `TCA_tot`, the plot highlights one of the fundamental properties of the profile likelihood: invariance under reparameterization. In comparison to the profiles without steady-state information, the new profiles are narrower, meaning that the parameters have smaller confidence intervals. This was to be expected because we have reduced the dimension of the parameter space by one.

**Figure 9:**
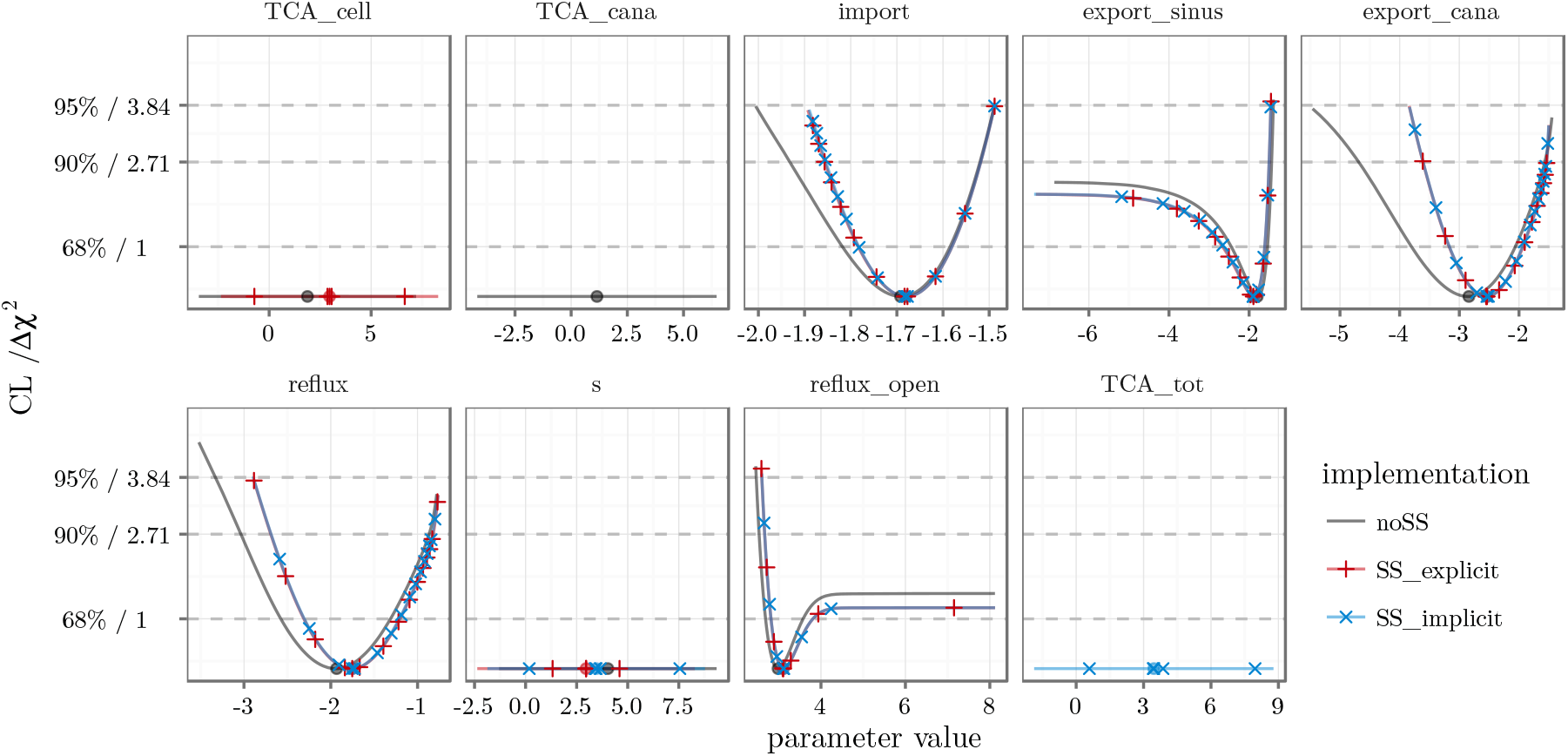
Parameter profiles for three different model implementations. The profile likelihood around the global optimum for the models without steady-state constraints, explicit steady-state constraints and implicit implementation of steady states is visualized by different colors. To illustrate that explicit (red) and implicit (blue) steady-state implementations yield the same result, the corresponding profiles are highlighted by red plus and blue cross signs, respectively.

### 4.8. Prediction uncertainty and validation profiles

Combining the steady-state constraint and two efflux experiments, one with closed canaliculi and the other with open canaliculi, we could fully identify the rate parameters `import`, `export_cana` and `reflux`. The amount parameter `TCA_tot` is fully coupled with the scaling parameter s such that both are structurally non-identifiable. The parameters `export_sinus` and `reflux_open` are practically non-identifiable since both parameters cannot be constrained to a finite interval with 95% confidence.

Next, we investigate the possibility to predict cellular amounts of TCA, `TCA_cell`, despite the non-identifiability of parameters. The amount of `TCA_cell` certainly depends on the total amount of TCA in the system. This total amount must be fixed in which case the parameters `TCA_cell`, `TCA_cana` and `s` become identifiable. The prediction uncertainty is assessed by a *prediction profile* which is computed based on a virtual data point for cellular TCA, measured at time point *t* = 41 under the “standard” condition. The **dMod** formulation reads as:

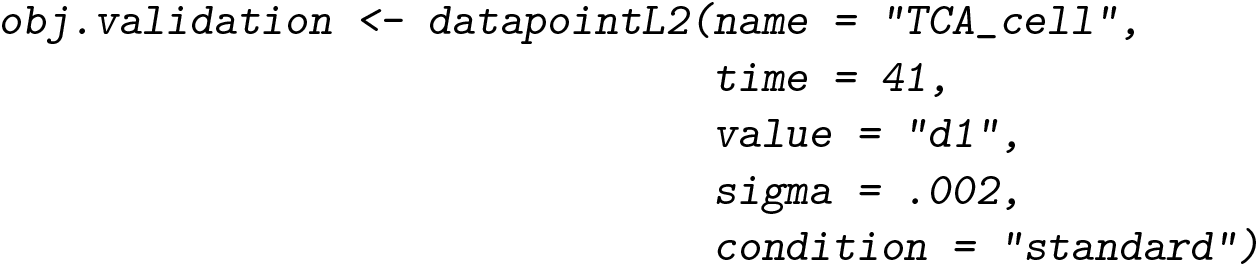

The uncertainty *σ* = 0.002 is set to a small value, i.e. below 1% of the prediction value. The `datapointL2()` command returns an objective function which evaluates the model prediction^1^ and computes the least-squares function of the virtual datapoint, returning derivatives for the data-point parameter d1. Its value “d1” is yet to be determined. By fitting the objective function obj together with obj.validation, “d1” equals the value of the `TCA_cell` at *t* = 41, as only then its contribution to the objective value is zero.

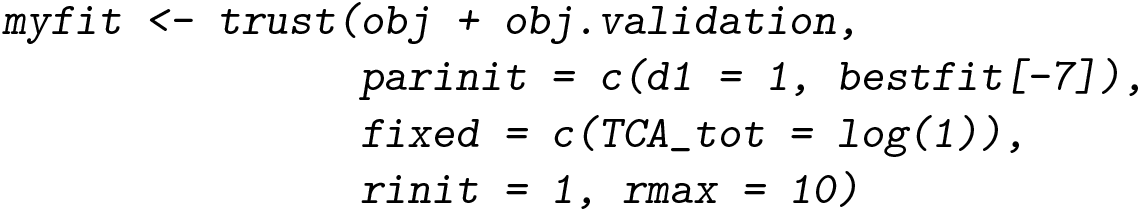

Using the derivative information provided by `datapointL2`, a prediction profile around `d1` is calculated.

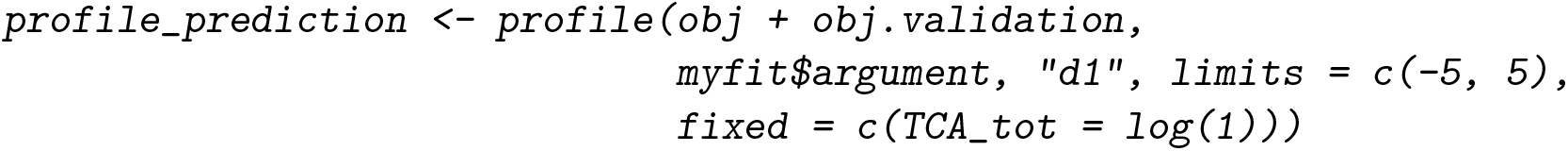

The result is shown in Figure 10A. The interpretation of such a prediction profile is, that a measurement yielding a value for the cellular TCA level at time point *t* = 41 outside of the interval [0.19, 0.21] does not conform to our model with 95% confidence.

**Figure 10:**
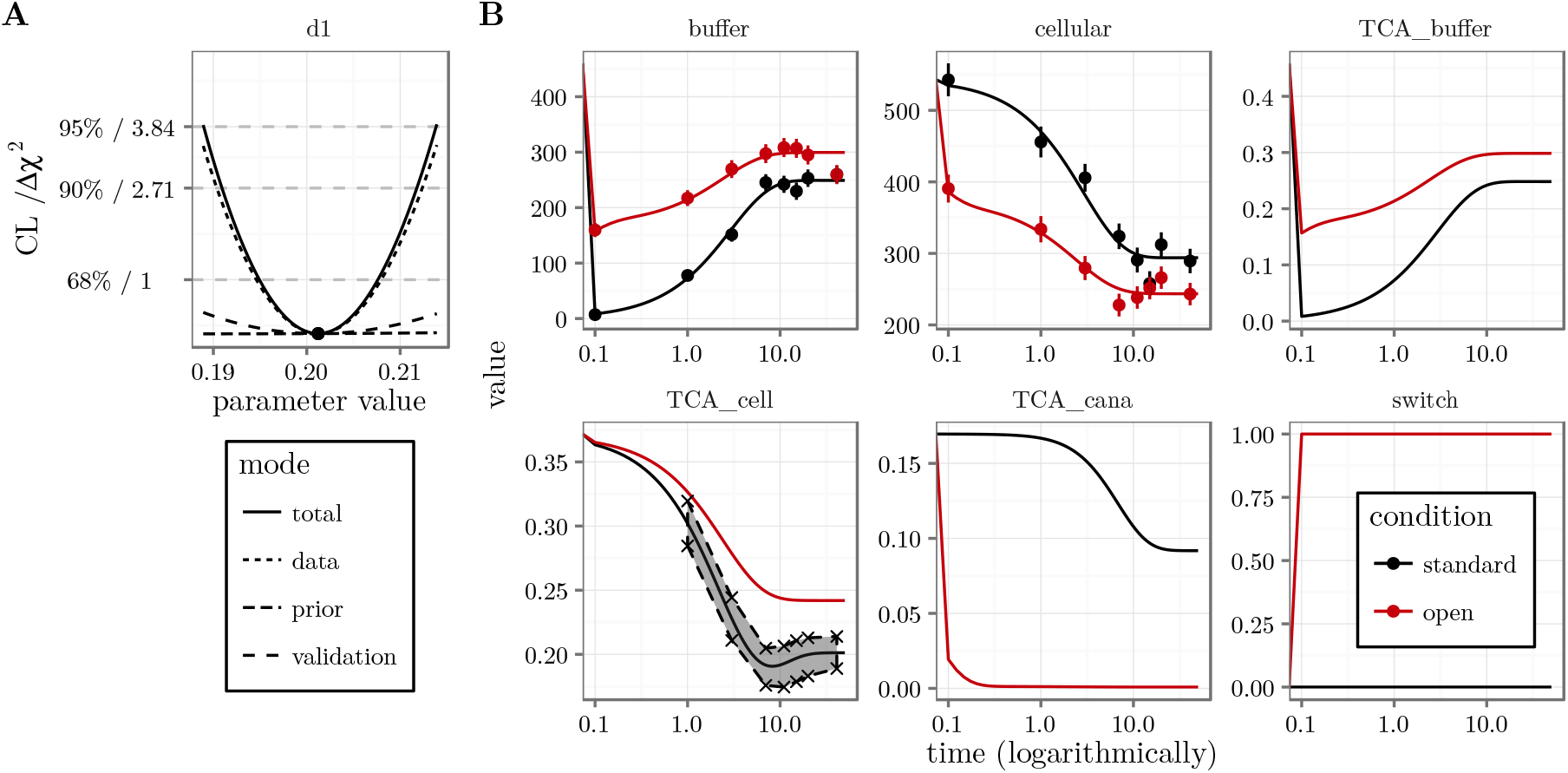
Validation profile and confidence bands for the model prediction. (A) The profile likelihood for the data point parameter d1 describing `TCA_cell` at time point *t* = 41 is shown. (B) Computing the data parameter profile for different time points yields 95% confidence bands on the prediction of `TCA_cell`.

More precisely, changing the data-point parameter d1, the model is quickly forced to follow the new data point. This is apparent from the “validation” contribution to the total objective value, Figure 10A, since it remains small with respect to the “data” contribution originating from all other data points. Forcing the model prediction to deviate more than 0.01 from the original one, the profile exceeds the 95% confidence threshold providing a confidence interval for the prediction itself. By calculating prediction profiles for several time points, a confidence band for the course of `TCA_cell` is constructed as shown in Figure 10B. The 95% confidence band is closed towards small and large amounts.

In summary, we find that the prediction of cellular TCA amounts is highly precise despite the non-identifiability of the `export_sinus` and `reflux_open` parameters.

## 5. Extensions of dMod

Computer algebra and symbolic tools are not part of R’s core functionality. In this section we illustrate two symbolic tools that are shiped with **dMod**, dealing with structural non-identifiability and steady-state constraints. They are implemented in Python and are interfaced via the **rPython** package.

### 5.1. Lie-group symmetry detection

In Section 4.6, profile likelihood computation showed the existence of both practically and structurally non-identifiable parameters. While practical non-identifiability arises from insufficient information in the data, structural non-identifiabilty is connected to Lie-group symmetries, i.e. transformations of the states and parameters

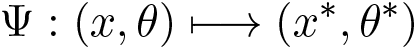

that preserve the model prediction of the observables:

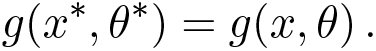

Based on (Merkt *et al.* 2015), the `symmetryDetection()` command outputs a list of available symmetry transformations. For example, the code

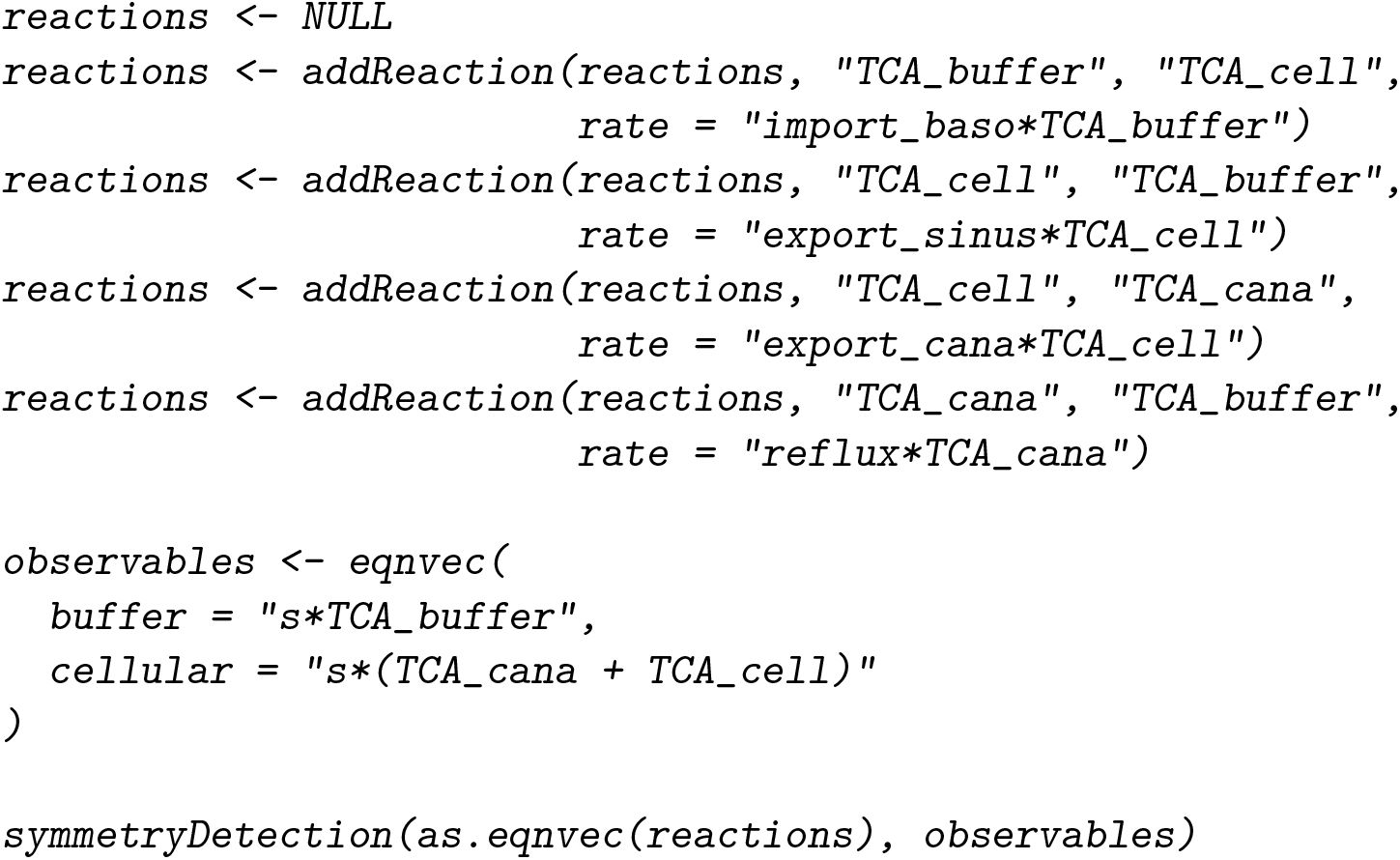

 returns the following output:

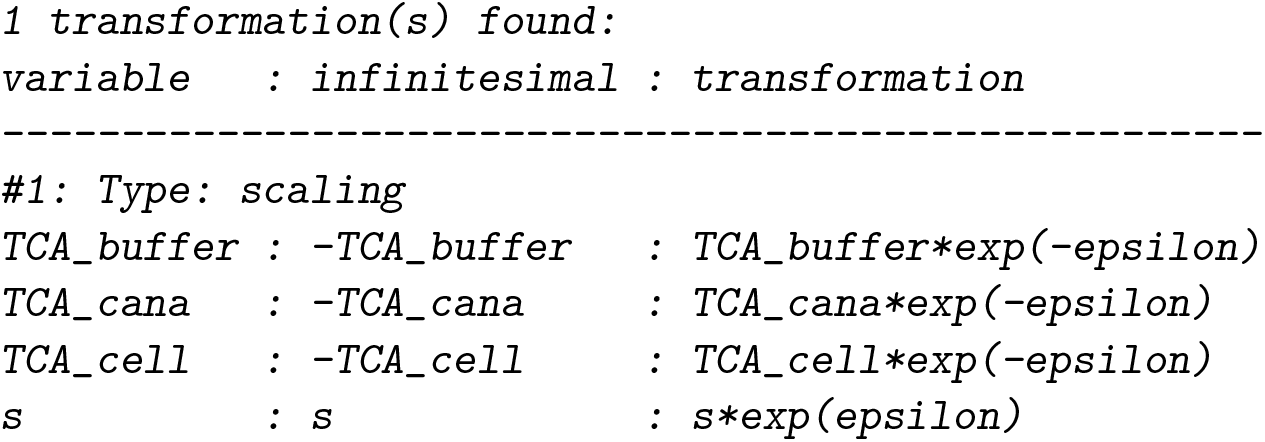

In agreement with the identifiability analysis by the profile-likelihood method, the parameters `s`, `TCA_buffer`, `TCA_cana` and `TCA_cell` are found to be non-identifiable due to a scaling symmetry. The corresponding scaling transformation, last column, leaves the observation invariant for any choice of epsilon. The parameter non-identifiability can be resolved choosing one representative from the orbit of the transformation. In our case, the scaling parameter could e.g. be fixed to 1.

### 5.2. Analytical steady-state constraints

While for the the present model, the steady-state could be explicitly calculated by hand, this might be much more challenging for models with a large number of states and parameters. For many of these models, the `steadyStates()` command outputs an analytical steady-state solution that can be incorporated in the model as an additional parameter transformation. Based on (Rosenblatt *et al.* 2016), the steady-state constraint is solved for a combination of state variables and kinetic parameters while positivity of the solution is ensured. As the paper states, the approach outperforms common methods of steady-state implementation with respect to reliability and performance of the optimization process. For our example, the code reads

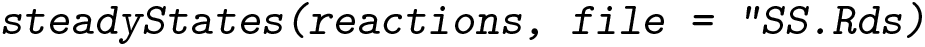

yielding the output

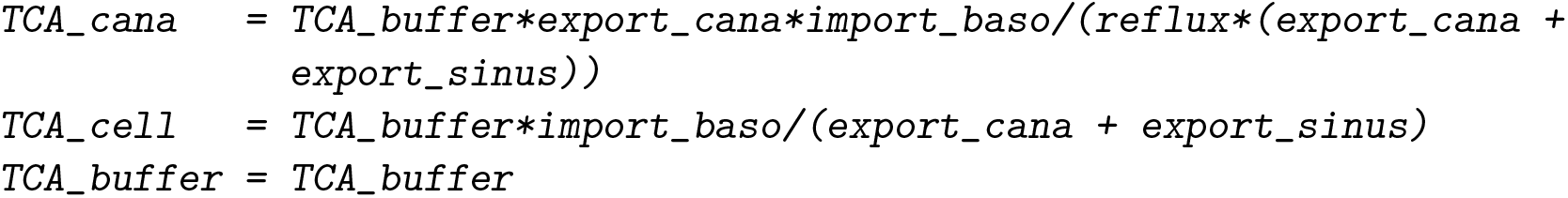

The solution is stored in an Rds file. After loading the file by readRDS(), the equations can be used when defining the parameter transformation.

## Footnotes

1 Several objective functions combined by the “+” operator share the same environment. Thus, the prediction computed by the first objective function can be evaluated by all other functions to come.

## References

Azzalini A (1996). Statistical Inference Based on the Likelihood, volume 68. CRC Press.

Bihorel S (2014). scaRabee: Optimization Toolkit for Pharmacokinetic-Pharmacodynamic Models. R package version 1.1-3, URL http://CRAN.R-project.org/package=scaRabee.

Hass H, Kreutz C, Timmer J, Kaschek D (2016). “Fast Integration-Based Prediction Bands for Ordinary Di erential Equation Models.” Bioinformatics, 32(8), 1204–1210.

Kreutz C, Raue A, Kaschek D, Timmer J (2013). “Pro le Likelihood in Systems Biology.” FEBS Journal, 280(11), 2564–2571.

Kreutz C, Raue A, Timmer J (2012). “Likelihood Based Observability Analysis and Confidence Intervals for Predictions of Dynamic Models.” BMC Systems Biology, 6(1), 120.

Maiwald T, Hass H, Steiert B, Vanlier J, Engesser R, Raue A, Kipkeew F, Bock HH, Kaschek D, Kreutz C, Timmer J (2016). “Driving the Model to Its Limit: Pro le Likelihood Based Model Reduction.” PloS ONE, 11(9), e0162366.

Merkt B, Timmer J, Kaschek D (2015). “Higher-Order Lie Symmetries in Identi ability and Predictability Analysis of Dynamic Models.” Physical Review E, 92(1), 012920.

Murphy SA, Van der Vaart AW (2000). “On Pro le Likelihood.” Journal of the American Statistical Association, 95(450), 449–465.

Press WH, Teukolsky SA, Vetterling WT, Flannery BP (1996). Numerical Recipes in C, volume 2. Cambridge University Press Cambridge.

Ranke J, Lindenberger K, Lehmann R (2016). mkin: Kinetic Evaluation of Chemical Degradation Data. R package version 0.9.44, URL http://CRAN.R-project.org/package=mkin.

Raue A, Kreutz C, Maiwald T, Bachmann J, Schilling M, Klingmüller U, Timmer J (2009). “Structural and Practical Identi ability Analysis of Partially Observed Dynamical Models by Exploiting the Pro le Likelihood.” Bioinformatics, 25(15), 1923–1929.

Raue A, Kreutz C, Maiwald T, Klingmüller U, Timmer J (2011). “Addressing Parameter Identi ability by Model-Based Experimentation.” IET Systems Biology, 5(2), 120–130.

Raue A, Kreutz C, Theis FJ, Timmer J (2013a). “Joining Forces of Bayesian and Frequentist Methodology: A Study for Inference in the Presence of Non-Identi ability.” Phil. Trans. R. Soc. A, 371(1984), 20110544.

Raue A, Schilling M, Bachmann J, Matteson A, Schelker M, Kaschek D, Hug S, Kreutz C, Harms BD, Theis FJ, Klingmüller U, Timmer J (2013b). “Lessons Learned from Quanti-tative Dynamical Modeling in Systems Biology.” PloS ONE, 8(9), e74335.

Rosenblatt M, Timmer J, Kaschek D (2016). “Customized Steady-State Constraints for Parameter Estimation in Non-Linear Ordinary Di erential Equation Models.” Frontiers in Cell and Developmental Biology, 4.

Soetaert K, Petzoldt T (2010). “Inverse Modelling, Sensitivity and Monte Carlo Analysis in R Using Package FME.” Journal of Statistical Software, 33(3), 1–28. URL http://www.jstatsoft.org/v33/i03/.

Soetaert K, Petzoldt T, Setzer RW (2010). “Solving Di erential Equations in R: Package deSolve.” Journal of Statistical Software, 33(9), 1–25. ISSN 1548-7660. URL http://www.jstatsoft.org/v33/i09.

Squire W, Trapp G (1998). “Using Complex Variables to Estimate Derivatives of Real Functions.” SIAM Review, 40(1), 110–112.

Tornoe CW (2012). nlmeODE: Non-Linear Mixed-effects Modelling in nlme Using Differential Equations. R package version 1.1, URL http://CRAN.R-project.org/package=nlmeODE.

Wright S, Nocedal J (1999). “Numerical Optimization.” Springer Science, 35, 67–68.

